# Combining rational design and continuous evolution on minimalist proteins that target DNA

**DOI:** 10.1101/2020.02.21.959445

**Authors:** Ichiro Inamoto, Inder Sheoran, Serban C. Popa, Montdher Hussain, Jumi A. Shin

## Abstract

We designed **MEF** to mimic the basic region/helix-loop-helix/leucine zipper (bHLHZ) domain of transcription factors Max and Myc, which bind with high DNA sequence specificity and affinity to the E-box motif (enhancer box, CACGTG). To make **MEF**, we started with our rationally designed ME47, a hybrid of the Max basic region and E47 HLH, that effectively inhibited tumor growth in a mouse model of breast cancer. ME47, however, displays propensity for instability and misfolding. We therefore sought to improve ME47’s structural and functional features. We used phage-assisted continuous evolution (PACE) to uncover “nonrational” changes to complement our rational design. PACE mutated Arg12 that contacts the DNA phosphodiester backbone. We would not have rationally made such a change, but this mutation improved ME47’s stability with little change in DNA-binding function. We mutated Cys29 to Ser and Ala in ME47’s HLH to eliminate undesired disulfide formation; these mutations reduced E-box binding activity. To compensate, we fused the designed FosW leucine zipper to ME47 to increase the dimerization interface and improve protein stability and E-box targeting activity. This “franken-protein” **MEF** comprises the Max basic region, E47 HLH, and FosW leucine zipper—plus mutations that arose during PACE and rational design—and is a tractable, reliable protein *in vivo* and *in vitro.* Compared with ME47, **MEF** gives three-fold stronger binding to E-box with four-fold increased specificity for E-box over nonspecific DNA. Generation of **MEF** demonstrates that combining rational design and continuous evolution can be a powerful tool for designing proteins with robust structure and strong DNA-binding function.

## INTRODUCTION

Transcription factors (TFs) often employ well-folded, stable α-helices to recognize and bind the DNA major groove with high sequence specificity and affinity.^1,2^ In addition, dimeric TFs also use α-helices for specific, high affinity protein-partner recognition: e.g., the <u>b</u>asic region/leucine <u>zip</u>per (bZIP), <u>b</u>asic region/<u>h</u>elix-loop-<u>h</u>elix (bHLH), and <u>b</u>asic region/<u>h</u>elix-loop-<u>h</u>elix/leucine <u>z</u>ipper (bHLHZ) are TF families that use α-helices for both protein-DNA and protein-protein interactions. Because TFs are highly modular in structure, the bZIP, bHLH, or bHLHZ domain can be expressed on its own—without any transcriptional activation domain or ligand-binding domain, for example—and still retain properly folded structure and specific DNA- and protein-binding functions. Despite the structural relatedness of these TF families, their α-helical protein dimerization interfaces are highly specific—whether homo- or heterodimeric partnering—and do not dimerize with members of other TF families.^3-5^

The modular, yet highly specific, nature of these protein motifs serves as the basis for our design of minimalist proteins that target specific DNA sequences. These compact domains can provide a practical tool for researchers to utilize toward exploring structure and function of larger transcription factors.^1,6^ These domains are also finding increased value in drug design; e.g., small proteins and peptidomimetic derivatives that target a particular TF can competitively inhibit and regulate the TF’s activities, including its protein-DNA and protein-protein interactions (reviewed in ref. 7). Omomyc, a bHLHZ protein that is a Myc derivative, has shown great promise as a “mini-protein” inhibitor of Myc that is undergoing patenting for anti-cancer therapeutic development.^8^ More and more, researchers are finding utility in employing not only rational design approaches, but also methods including phage display and library studies that are potentially hastening the development of proteins as drugs.^9-11^

When we embark on design of a minimalist protein, we first employ a rational design approach, wherein we fuse subdomains from the bZIP, bHLH, and bHLHZ motifs to make hybrid proteins that rival the high binding affinity and DNA sequence specificity displayed by native TFs for their DNA target sites. Rational design refers to our use of literature, modeling, and knowledge to generate unique proteins with desired traits: in our case, binding to a specific DNA site, tractable size, and stable structure. ME47, previously termed “Max-E47,” is an example of a rationally designed TF.^5^ ME47 comprises 66 amino acids (aa) and is a hybrid of the DNA-binding basic region from Max, a bHLHZ protein, and the HLH dimerization domain from E47, a bHLH protein.^5,12,13^ Another protein, MFW (84 aa, previously termed “MaxFosW”^14^), is a modified Max bHLHZ in which the native Max leucine zipper (LZ) is replaced by FosW,^15,16^ which is a designed LZ variant based on the bZIP protein c-Fos.^17-19^ Interestingly, the engineered FosW homodimerizes, while the native c-Fos LZ does not. Our rational design of minimalist proteins (<100 aa) depends on knowledge of protein structure, e.g., the Max bHLHZ domain (92 aa) has crystal structures.^20-22^ *De novo* design of proteins in this size range is arguably beyond current state of the art, although Baker has achieved significant success with ∼40 aa proteins using a massively parallel approach that integrated large-scale computation, yeast display screening, and next-generation sequencing.^23^

Although subdomain swapping within the same TF family has been successful,^24-27^ such swaps *between* different TF families are uncommon. To our knowledge, we are one of two groups to report design of *inter*-family hybrid proteins: these include our hybrids of the bHLH modules of bHLH/PAS proteins AhR and Arnt fused to LZs from bZIP proteins C/EBP and Fos,^5,14,28,29^ and Chapman-Smith and Whitelaw generated a hybrid comprising the bHLH from Arnt and LZ from Max.^30^. Our fusion proteins target the E-box DNA site (<u>e</u>nhancer box, 5’-CACGTG) that is bound by bHLHZ TFs Max and Myc,^20-22,31-34^ which are involved in numerous normal cellular processes such as cell cycle regulation and differentiation, but whose abnormal activities are associated with over half of all human cancers.^35^

More recently, we have worked towards incorporating “nonrational” design to generate minimalist proteins with improved E-box targeting activity. Initially, we attempted to improve ME47’s E-box binding properties by extensively using library screening in yeast or bacterial one-hybrid assays (Y1H or B1H).^36-38^ However, our experiments were burdened with numerous false positive signals and difficulty incorporating large, unbiased libraries,^12^ which are challenges common to *in vivo* screening and selection assays.^39,40^ To circumvent these problems, we turned to evolution-based methods, such as continuous evolution in PACE (<u>p</u>hage-<u>a</u>ssisted <u>c</u>ontinuous <u>e</u>volution).^41^ In PACE, only genes—and mutations to those genes—that confer a survival advantage are propagated within the population of modified M13 phage. Since mutations are constantly introduced at every generation by an inducible error-prone DNA polymerase, PACE is now termed “continuous” evolution, as opposed to approaches like phage display that introduce mutations at set points during the experiment (reviewed in ref. 42). Continual mutagenesis coupled with selective pressure is the hallmark difference between evolution vs. screening/selection, in which the size and quality (e.g., lack of bias in the sequences) of the starting library can significantly impact the experimental outcome.

ME47 has performed well in various *in vivo* and *in vitro* assays that demonstrated its α-helical structure and strong binding to the E-box site. In addition, expression of ME47 in an established tumor xenograft of a triple negative basal breast cancer inhibited tumor growth in a mouse model by blocking Myc’s transcriptional activity from the E-box.^43^ Thus, ME47 has proven to have excellent functional activity as an E-box binder, even as a competitive inhibitor of Myc/Max binding to the E-box in a mouse model of human cancer. However, we find ME47 displays instability stemming from misfolded structure, and we have long observed that bHLH and bHLHZ domains can have propensity for misfolding and aggregation.^5,12,14,28,29^ Likewise, other groups have encountered the same problems with the Max bHLHZ and found it difficult to manipulate when measuring the binding of the Max bHLHZ to the E-box.^44-46^ These domains have been found to contain significant structural disorder, and the Max and Myc bHLHZ domains were recently classified as intrinsically disordered proteins.^47-49^

We therefore decided to improve ME47’s structure and function through a combination of rational design and continuous evolution to make it more experimentally tractable. The resultant protein is **MEF** (89 aa), in which we turned a bHLH protein (ME47 is a bHLH domain^13^) into a bHLH**Z**-type of structure by appending the non-native LZ FosW to its C-terminus (Fig. 1).^15^ **MEF** is a hybrid protein comprising modules from three different protein families and is the most complex minimalist hybrid protein we have generated. **MEF** comprises the basic region from Max bHLHZ,^20^ the four-helix bundle dimerization element from E47 bHLH,^50^ and the FosW LZ: hence, **MEF** was designed to retain the E-box binding function exhibited by the Max basic region and maintain a dimeric bHLHZ structure that preferentially homodimerizes via the unnatural interface provided by the E47 HLH and FosW LZ. We used the *in vivo* B1H assay to assess protein-DNA complexes in live *Escherichia coli* (*E. coli*) cells. We used *in vitro* methods quantitative electrophoretic mobility shift assay (EMSA) to assess binding affinities of protein-DNA complexes and circular dichroism (CD) to assess protein secondary structure in order to characterize our “franken-protein” **MEF** and variants that maintain the intended α-helical structure and high-affinity E-box binding function. Our work demonstrates the clear advantages of combining our rational design know-how with the unpredicted, yet beneficial, mutations uncovered via PACE to design the more structurally consistent and robust minimalist protein **MEF** with improved E-box binding function.

**Figure 1.**
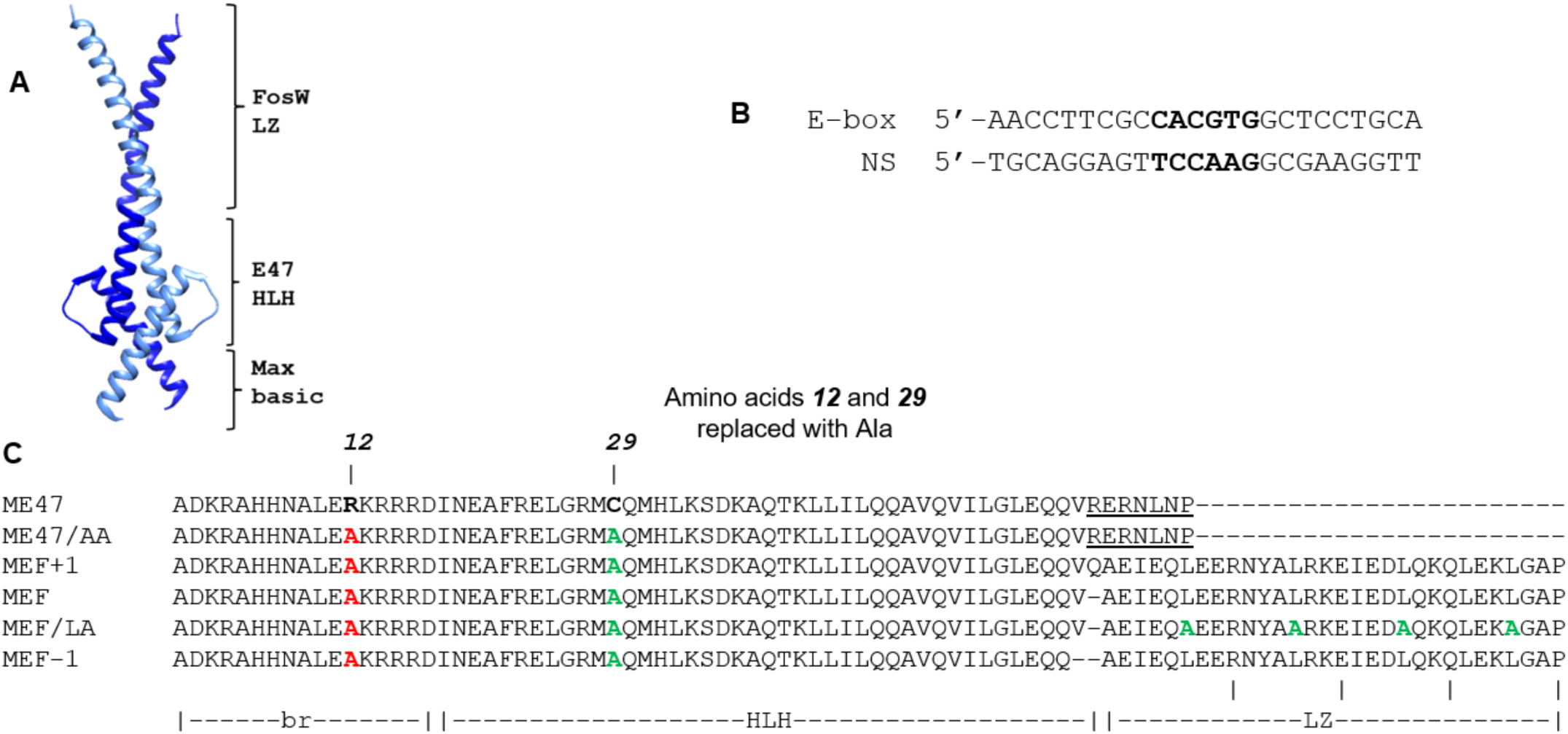
A. Representative MEF structure. The ME47 (PDB:3U5V^13^) and FosW (PDB: 5FV8^51^) crystal structures were merged to illustrate the intended structure of the MEF dimer; monomers highlighted in light and dark blue (Chimera v. 1.11.2^52^). **B. DNA sequences used in reporter constructs for B1H, CD, and EMSA**. E-box (CACGTG) and NS (TCCAAG) sequences are bolded. **C. Sequence alignment.** Residues in red were mutations that arose from PACE; residues in green were rationally designed. Underlined residues of ME47 sequence were replaced with the FosW LZ, which was appended right after residues QQV in ME47 to place the FosW LZ in proper register (see Fig. S13 for illustrations of in- and out-of-register coiled coil structures). Dashes indicate gaps in the protein sequence that maintain alignment; br, basic region; HLH, helix-loop-helix; LZ leucine zipper.

## RESULTS & DISCUSSION

### Continuous evolution on ME47 yielded an unpredicted mutation at Arg12

Minimalist ME47 comprises 66 amino acids, compared to the bHLHZ domains of Max and Myc that comprise 92 amino acids with their additional length from the leucine zippers that are necessary for dimerization. ME47 utilizes E47’s HLH for homodimerization; as a bHLH transcription factor, E47 natively and effectively dimerizes via the HLH without the aid of a LZ. Although ME47 demonstrates strong and specific binding of the E-box rivaling that of native TFs, it has propensity to misfold, whether during CD measurements showing unusually low α-helicity or *K*_d_ measurements via fluorescence anisotropy or EMSA that show weak or nonspecific binding of ME47 to the E-box. Numerous attempts to improve ME47’s structure and function by extensive *in vitro* selection from random libraries and *in vivo* screening in the Y1H and B1H were unsuccessful.^12^

We therefore embarked on directed evolution of toward improving ME47; in particular, we encountered the PACE methodology,^41^ and we hypothesized that continuous evolution combined with our rational design approach could be used together to improve ME47. PACE utilizes infection-propagation cycles of M13 bacteriophage on its *E. coli* host as the medium for continuous evolution. The ability of M13 phage to propagate functional offspring is linked to the activity of the gene of interest to be evolved. This is done by deleting gIII (coding for pIII, an essential structural protein) from the phage genome, thereby rendering the phage incapable of propagating by itself. A reporter gene is constructed where gIII expression is controlled by the protein of interest, and the reporter is added to the system on a host-borne plasmid. Phage propagation correlates with the activity of the protein of interest. Thus, mutations in the protein that enhance activity will confer survival advantage for the phage, while detrimental mutations will hinder the phage: this is the basis of evolution in PACE. Mutations can be introduced every generation into the coding sequences of proteins via an inducible error-prone DNA polymerase. Our re-designed PACE system exerts selective pressure such that enhanced E-box binders continue to subsequent generations.^53^

To evolve ME47 in PACE, we constructed an E-box-regulated gIII reporter circuit by borrowing from the bacterial one-hybrid (B1H) system.^37,38,54^ We began by optimizing the E-box regulated reporter gene in the B1H system (Fig. S1; the spacer is the linker between the 3’ end of the E-box and beginning of the -35 position of the weak lac promoter; refer to Fig. S2 for more details).^53,55^ After extensive exploration of spacer lengths ranging from 7 to 25 base pairs (bp), we established that the -7GC E-box reporter was appropriate for use with ME47 in the context of PACE, and that this reporter was only activated by ME47 and not by any endogenous cellular proteins (Figs. 1, S2, S3, S4; “7” refers to spacer length in bp, “GC” refers to flanking sequence variants 2 bp outside of the E-box, Fig. 1B). Henceforth, this -7GC E-box is termed as the “E-box.”

ME47 was evolved on the E-box reporter gene for seven days. By day three, ME47 with mutations altering Arg12 to Cys were present in >50% of the population (Fig. S5). After five days, all of the phage population sequenced contained contained either Cys12 or Ser12. These mutants are ME47/CC and ME47/SC, respectively (Table 1; refer to Table 1 and the Experimental Section for explanation of protein nomenclature).

**Table 1.**
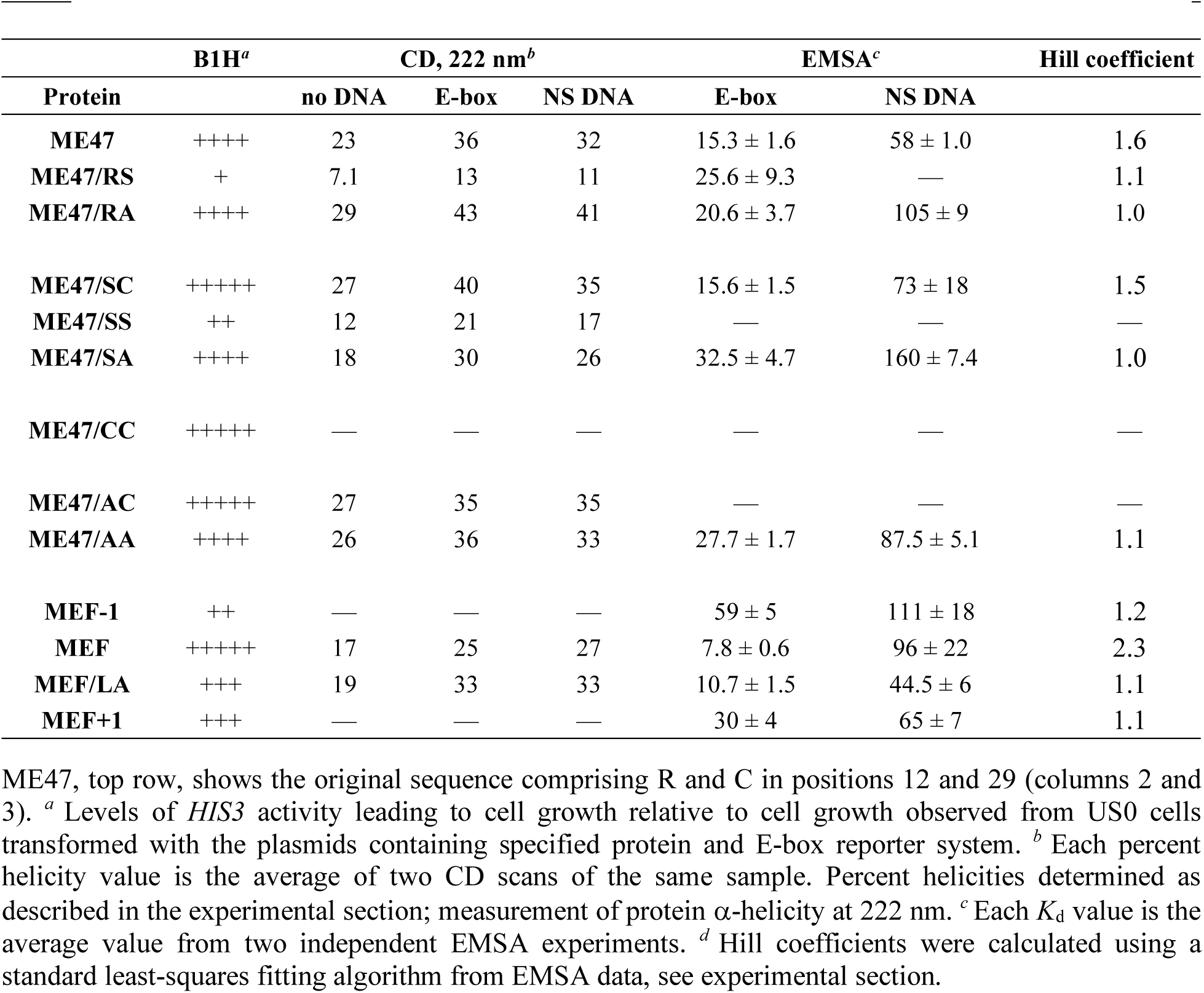
Protein-DNA complexes. B1H,^*a*^ CD,^*b*^ *K*_d_ values from EMSA,^*c*^ Hill coefficients.^*d*^

### Mutation of Arg12 to an uncharged residue unexpectedly improved ME47’s E-box binding function

We were excited to uncover this Arg12 mutation, but initially, we were skeptical of the validity of this mutation given that in the Max crystal structure, Arg12 makes a Coulombic interaction with the DNA phosphodiester backbone at the highlighted central G in the core E-box motif CACGTG (Fig. S6).^20^ Significantly, protein contacts with nucleotides at this position determine whether TFs interact with the canonical Class A E-box (CACGTG) or non-canonical Class B E-box (CAGCTG).^56^ As such, we were mystified as to why mutations at this position outcompeted the original ME47 in PACE. We therefore validated the Arg12 mutations outside of the context of PACE, where phage-host interactions may confound our analysis. To do so, we re-constructed ME47/SC and ME47/CC independently from the PACE system and assessed them in the B1H and EMSA (Table 1). In the B1H, ME47/SC and ME47/CC outperformed ME47, producing a much higher transcriptional activation signal at the E-box (Fig. 2; note that the signal reporting on protein-DNA binding in the B1H is *not* linear, and that each column shows a 10-fold dilution difference in cell growth). Interestingly, the mutants did not show improved binding to E-box when measured by EMSA. B1H and EMSA on the nonspecific DNA control sequence (NS DNA) showed no signal, thereby confirming that these mutants did not interact nonspecifically with DNA (Fig. S7A).

**Figure 2.**
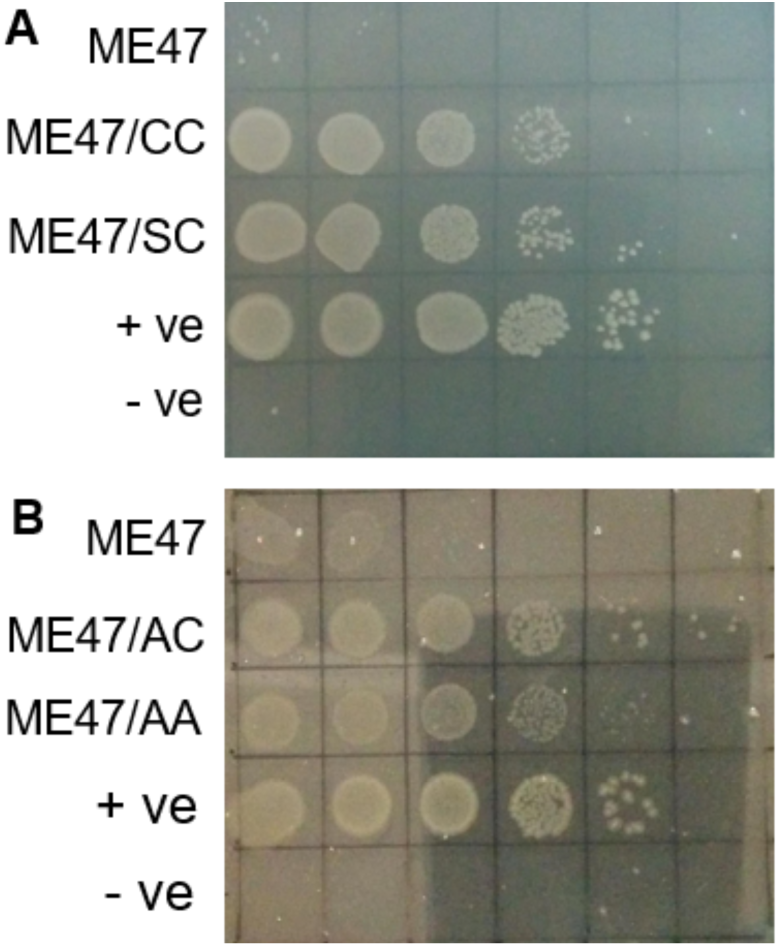
B1H results for Arg12 mutants of ME47. **A.** E-box-responsive transcriptional activation of ME47 Arg12 mutants derived from PACE, plated at a 10-fold serial dilutions (10^−1^-10^−6^ from left to right) after initially diluting cells to OD_600_ ∼0.1 on a 20 mM 3-AT plate in the B1H assay. Both ME47/CC and ME47/SC, which are the Arg12Cys and Arg12Ser variants of ME47, produce signals that are much stronger relative to that from ME47. The positive control (**+**) comprises transcription factor Zif268 with its cognate DNA site in lieu of E-box; the negative control (**-**) comprises Zif268 with an unchanged pH3U3 vector that lacks the Zif268 cognate site upstream of the *HIS3* reporter.^37^ **B.** E-box-responsive transcriptional activation of ME47/AC (Arg12Ala mutant) that was rationally designed based on PACE mutants in **A**, plated as above. The ME47/AC mutant produced signal that is much stronger than signal from ME47. **ME47/AA** is slightly weaker than ME47/AC *in vivo*, but outperformed it in terms of reliability *in vitro*, which led to it being the final iteration of ME47 design. Note that ME47 shows much weaker transcriptional activation from the E-box compared with any of the mutants.

Surprisingly, these Arg12 mutants that arose from PACE performed markedly better than original ME47 in the B1H assay—despite apparently losing a Coulombic contact with the cognate E-box—thereby validating that the Arg12 mutations did indeed improve the activity of ME47. These observations led us to explore more mutations at this position; because we wished to avoid complications from disulfide formation between Cys residues, we did not pursue Cys in position 12. In our previous work on the rational design of bZIP proteins, we demonstrated that as many as 24 out of 27 residues in the basic region could be Ala, which is the best α-helix former and stabilizer,^57,58^ despite that basic regions are enriched with positively charged residues that interact with DNA. These Ala mutations had no impact on the DNA-binding properties of the bZIP protein, likely due to compensatory effects in the protein-DNA complex: increased α-helicity and protein stability can compensate for enthalpic losses from reduced Coulombic interactions, for example.^27,59^ These previous results inspired us to substitute Ala into position 12 to generate ME47/AC, which showed excellent E-box binding affinity and specificity in the B1H (Fig. 2B, Table 1).

We then created ME47 variants with Arg12 mutated to Thr or Val and tested them in the B1H assay (Fig. S8); these mutations would test how size of the amino acid side chain and even hydrophobicity from an additional methyl group impact function, as well as the importance of the hydroxyl/sulfhydryl moiety. The introduction of Val12 improved binding in the B1H, albeit not as large an improvement as with the Ser12 and Thr12 mutants. Taken together, the results from these mutants at position 12 suggest that smaller residues—and not the native, bulkier Arg from the Max basic region—are beneficial to ME47 activity; as ME47 is a franken-protein, it is possible that these smaller residues allow the protein to assume a conformation that is more conducive to interacting with the E-box.

### Arg12 mutations may improve structural stability by a mechanism unique to ME47

The Arg12 mutations improved the effectiveness of ME47 in PACE and B1H, but not in EMSA. Both PACE and B1H are *in vivo* experiments where the output of each experiment is the sum of assorted parameters associated with the particular live host cell, whereas in EMSA, the results can more directly correlate with *in vitro* protein-DNA binding affinity, as there are no confounding effects from unrelated cellular processes, for example. This led us to hypothesize that by mutating Arg12, PACE may have optimized other features of ME47, aside from binding specificity and affinity. One such feature could be protein robustness, which refers to stability and foldedness that could confer a survival advantage during an evolution that runs over many days.

To investigate if PACE can allow proteins to improve features like robustness, we conducted a “competition PACE,” where proteins compete for survival with no induction of mutagenesis: hence, survival of each protein was based purely on its activity (outside of the rare occurrence of a random background mutation). This competition started with equal amounts of phage expressing ME47, ME47/SC, ME47/CC, and MFW,^14^ which is a bHLHZ protein with ∼15-fold weaker affinity for the E-box than ME47 (*K*_d_ 240 nM vs. 15 nM, respectively). To our surprise, MFW dominated the culture by day 3, despite its markedly weaker affinity for E-box DNA. Hence, MFW must have used features other than binding affinity/specificity to outcompete the other proteins for dominance. We surmised that this was due to MFW’s structural stability, allowing it to persist in the host cell to contribute toward reporter gene expression, in comparison to the less stable ME47 and variants; Western analysis shows that MFW actually expresses less than the ME47 variants, so MFW dominating the ME47 variants in competition PACE is not due to high expression levels of MFW (Fig. S9). Interestingly, we have always found MFW to be a consistently folded, reliable “workhorse” protein in our studies. The observation that MFW dominated the competition PACE despite its weaker E-box binding affinity aligns with the results above where ME47’s Arg12 mutants outperformed ME47 in *in vivo* assays B1H and PACE, but not in EMSA. The ME47/SC and ME47/CC mutants may have dominated over ME47 in PACE, because Ser and Cys are better α-helix stabilizers than Arg,^57,58^ thereby leading to more structurally robust proteins, and not because those mutations directly improve E-box binding activity. Aside from binding function, PACE can apparently enhance structural and thermodynamic stability. Indeed, in their original PACE paper, Esvelt *et al*. noted that evolved improvements in activity were higher in cells than in *in vitro* studies, indicating that these improvements were specific to the context of the bacterial cell environment in which they evolved.^41^

### Testing transferrability of Arg12 mutations to MFW

Given these results with the Arg12 mutants of ME47, we explored whether the same mutation could be transferred to MFW. As mentioned earlier, MFW has the same Max basic region as ME47, but the HLH element is that of Max, not E47; MFW also has the FosW LZ and is a bHLHZ domain, while ME47 is bHLH. Thus, MFW should be structurally similar to native Max bHLHZ, whose Arg12 contact with the DNA phosphodiester backbone would be expected to contribute to DNA-binding affinity. In the B1H, MFW with Arg12 mutated to Ser showed significantly *reduced* transcriptional activation from the E-box (Fig. S10), while ME47 with Arg12 mutated to Ser, ME47/SC, showed the opposite (Fig. 2A). MFW with Arg12 mutated to Ala had comparable, if slightly improved, activity (Fig. S10), while ME47 with Ala12, ME47/AC, showed markedly increased transcriptional activation (Fig. 2B). These MFW mutants showed that the Arg12 mutation obtained for ME47 was not altogether transferrable to MFW, except that interestingly, Arg12 could be replaced by Ala and still maintain good activity, indicating that this Coulombic interaction is not important for both ME47 and MFW binding to E-box.

In ME47, the junction for fusion of the Max basic region and E47 HLH domain is just 3 aa from Arg12; thus, Arg12 may not behave the same in ME47 as it does in Max and MFW. We found that PACE could uncover mutation(s) synergistic to our rational design, in particular, mutations advantageous to creating robust proteins that remained active over days of evolution. We also observed that PACE on ME47 showed remarkable sensitivity to the DNA target sequence. ME47 has higher affinity for the E-box with AT bp in the -2 bp flanking position than to the GC-flanked E-box in the B1H (Fig. S2), even though the AT/GC flanks are at -2 bp positions *outside the core E-box*. Interestingly, mutants ME47/SC and ME47/CC lost the ability to distinguish between the AT and GC E-boxes and strongly activate transcription from both targets in the B1H (Table S1). This indicates evolution toward the intended target, as the GC E-box (i.e., the “E-box”) was used in PACE, as above. In contrast, EMSA did not show such distinction, as *K*_d_ values of all proteins to AT and GC E-boxes are similar (Table S1). Possibly, differences between the *in vivo* bacterial environment in PACE and B1H and the *in vitro* EMSA buffer may influence protein structural features that affect DNA-binding function.

We decided to move forward with ME47/AC, which showed excellent E-box activity in the B1H, but only modest activity in EMSA (Table 1). However, we still observed some intractability *in vitro* for the various Arg12 mutants, which necessitated further refinement of the protein scaffold.

### Replacement of Cys29 in the HLH dimerization unit of ME47 to Ser or Ala

ME47 possesses Cys29 in Helix 1 of the E47 HLH domain.^5^ Because we wanted to eliminate any possibility of Cys thiol side-chain reactivity, we made a version of ME47 with Ser29. Such a construct should behave virtually the same as the original, as Ser is structurally similar to Cys. Our ME47 crystal structure shows that the Cys29 side chain faces into the four-helix bundle, and thus, Ser29 should mimic this activity (Fig. 3).^13^ Despite these predictions, the Ser29 version (ME47/RS) was highly unreliable in all experiments (Table 1). ME47/RS displayed poor α-helicity by CD (7-11%) whether in the presence or absence of E-box DNA, and weak/modest E-box activity in both the B1H and quantitative EMSA (*K*_d_ 26 nM). Because Ser is routinely used to replace Cys when no disulfide is formed, we were perplexed by these results and decided to make the ME47/SS mutant, where we combined the Ser12 mutation uncovered by PACE with Ser29. This double mutant also performed poorly, particularly in the B1H, and displayed fairly weak helical structure by CD (Fig. S7B, Table 1). Possibly, the E47 HLH evolved to utilize the Cys29 to maintain the hydrophobic packing needed for dimerization, and introduction of the more polar Ser disrupts the hydrophobic interface, thereby weakening dimerization and diminishing overall E-box-binding function. We concluded that Ser29 was not a functional replacement for Cys29, at least in ME47.

**Figure 3.**
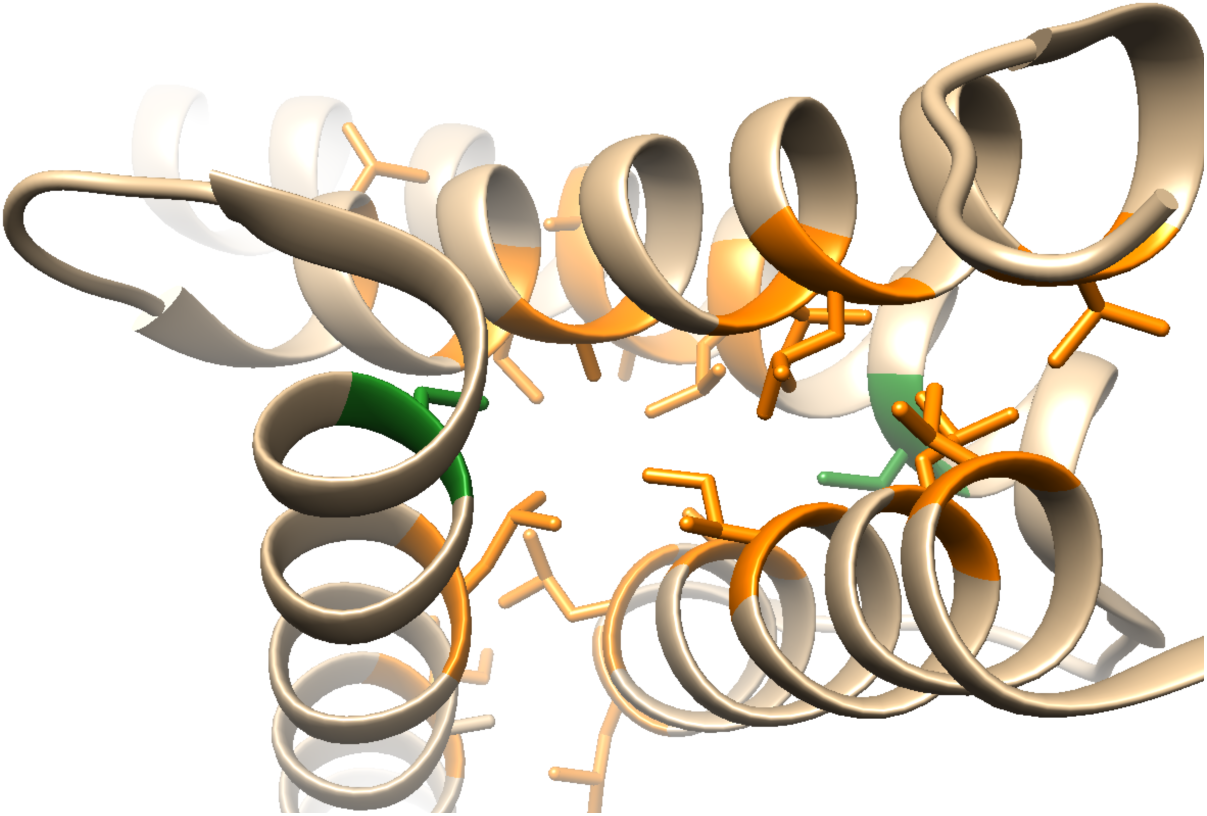
ME47 dimerization interface. The dimerization of ME47 occurs via hydrophobic packing of small, nonpolar residues (orange) found in Helices 1 and 2 (light brown, Chimera v. 1.11.2^52^). Cys29 residues (green) found at the ends of Helices 1 face inward toward the hydrophobic core. Mutating Cys29 to Ala greatly improved protein tractability in *in vitro* assays. In our ME47 crystal structure,^13^ the sulfur atoms in the Cys29 residues from the two monomers are 10.6 Å away from one another; given that typical disulfide distances are 2-3 Å,^60^ we did not expect Cys dimerization to occur.

Looking again at the ME47 crystal structure,^13^ we decided to substitute a small, hydrophobic residue in place of Cys29. We surmised that Ala29 would serve the same functionality that allows hydrophobic packing between the two monomers, and its smaller sidechain might reduce steric interference at the protein interface. Moreover, Ala29 would assist in making Helix 1 a more stable helical structure.^57,58^ Therefore, as in the case of using Ala12 in place of Arg12 in the previous section, we surmised that Ala29 may allow us to remove Cys29 with minimal negative impact on ME47. ME47/RA, which contains Ala29, is strongly helical by CD and shows superb binding to the E-box (*K*_d_ 7 nM) and much weaker nonspecific binding. But we note that in the B1H, the Ala29 mutant still is not as good at transcriptional activation from the E-box as original ME47, which contains Cys29 (Fig. S7B). Interestingly, Benezra made Ser and Ala mutants at the homologous Cys570 in Helix 1 of E12, which is an isoform of E47 (E12 also called E2A).^61^ This Cys570 was essential for homodimer stability and binding to the E-box, as the two Cys residues in the homodimer formed a covalent disulfide bond under certain cellular and temperature conditions; the Ser and Ala mutants at Cys570 of E12 showed significantly reduced homodimer formation and no ability for E-box binding. Similar to E12, our Cys29-containing mutants ME47/SC, CC, and AC all display very strong transcriptional activation from the E-box in the B1H, while the Ser29 and Ala29 mutants show greatly and slightly reduced activity, respectively.

We initially discounted that the two Cys29 residues in ME47 could form a disulfide linkage; our ME47 crystal structure shows that the two sulfur atoms are over 10 Å apart,^13^ which is well beyond the length of a disulfide bond (2.3 Å.^60^) Additionally, Hill coefficient analysis of two Cys29-containing proteins, ME47 and ME47/SC, give values 1.6 and 1.5, respectively, indicating that these proteins exhibit both cooperative dimer as well as noncooperative monomers binding to E-box: i.e., two monomers independently bound to each E-box half-site (cooperativity and Hill coefficients discussed in Supporting Information, Fig. S11). These Hill coefficients of ∼1.5 indicate a stronger possibility of cooperative dimeric binding than other values in Table 1 that show no cooperativity (Hill coefficients 1.0-1.1, Table 1). However, proteins are dynamic molecules, and their movements could facilitate a disulfide linkage in ME47 and ME47/SC. In particular, our B1H data showed that mutation of ME47’s Cys29 reduced transcriptional activation from the E-box, indicating that possibly the bacterial environment may foster the oxidizing conditions that promote Cys disulfide linkage. Regardless, as we wished to avoid Cys residues, we opted to pursue Ala29-containing mutants.

### Combination of nonrational PACE and rational design yielded ME47/AA

We next sought to combine the Arg12 and Cys29 mutations to generate the double mutant **ME47/AA** that contains both Ala12 and Ala29, though other permutations were also explored (Figs. 2B, Table 1). Although **ME47/AA** is highly tractable and predictable to use in all experiments, whether *in vivo* or *in vitro*, it is not a particularly strong E-box binder by the metrics shown in Table 1: e.g., the Cys29-containing mutants generally show the highest levels of transcriptional activation from the E-box in the B1H, followed by the A29-containing mutants of ME47. The *K*_d_ value for **ME47/AA** binding to the E-box (27 nM) is modest compared with other ME47 mutants with *K*_d_ values ∼15-17 nM, but CD does show that **ME47/AA** contains comparatively high α-helical secondary structure.

Because **ME47/AA** gives reliable and reproducible results, we decided to continue with this variant. However, we ultimately wanted a protein that is easy to work with *in vitro* without sacrificing E-box binding affinity and specificity, which brought us back to the drawing board to consider other changes we could make to the **ME47/AA** scaffold. Looking at the Hill coefficients (Table 1), we observed that **ME47/AA** appears to bind to the E-box as two monomers binding in the absence of cooperativity to the E-box. ME47 and ME47/SC—both containing Cys29—are the only ME47 mutants displaying Hill coefficients of 1.6 and 1.5, respectively, suggesting some cooperativity during E-box binding. This reminds us of similar mutants of E12 discussed above, where Cys residues in the HLH interface were involved in protein dimerization. Although our Ala29 mutation can contribute to helical secondary structure and hydrophobic packing of the HLH, these contributions do not fully compensate for removal of Cys29. Likewise, Benezra found that alanine’s hydrophobicity in his Cys570Ala mutant was insufficient to stabilize E12 homodimerization, although this mutant was still capable of heterodimerizing with MyoD.^61^ Thus, we rationalized that we must compensate for the Ala29 mutation in **ME47/AA** to design a protein with improved homodimerization ability that would translate into stronger E-box binding activity.

### Fusion of the FosW leucine zipper to ME47/AA to improve stability

Although our goal is to design *minimalist* proteins that target the E-box, we decided to append a leucine zipper to the **ME47/AA** bHLH to generate the bHLHZ domain. The LZ should assist in protein homodimerization—contributing to both partner specificity and binding affinity at the protein-protein interface—and thermodynamic stabilization of overall protein α-helical foldedness and structure: these advantages should compensate for enlarging our minimalist design. Both the bHLHZ and bHLH/PAS (Per-Arnt-Sim domain) transcription factor motifs utilize secondary dimerization elements (LZ, PAS) that contribute to protein dimerization and partner selection, although the bHLH family accomplishes these tasks with only one dimerization element (HLH).

We reasoned that a LZ secondary dimerization element could improve **ME47/AA**’s robustness and E-box-binding function. Previously, we used the LZ from bZIP transcription factor C/EBP to generate Max bHLHZ hybrids containing the C/EBP LZ in place of the native LZ, as well as swapping out the entire PAS element (330 aa) of the Arnt bHLH/PAS domain with the C/EBP leucine zipper (29 aa).^12,28^ Our newer hybrid proteins comprising the LZ of FosW,^15,16^ including both Max and Arnt derivatives with the FosW LZ,^14,29^ are more tractable and display better DNA-binding function than those hybrids with the C/EBP LZ, so we chose to append FosW to **ME47/AA**. The native c-Fos LZ heterodimerizes with partner c-Jun and does not homodimerize,^17-19^ but the re-designed FosW can homodimerize and heterodimerize. Notably, two Lys residues in the native c-Fos LZ coiled coil that prevent homodimerization were replaced with Asn and Ile.^15,16^ We also considered that a benefit of adding the FosW LZ is that it is a non-native designed construct, and it is based on c-Fos, a bZIP protein that will not dimerize with proteins outside of the bZIP family. Thus, the dimerization interface comprising the E47 HLH and FosW LZ should only homodimerize and not fortuitously partner with an endogenous protein in any type of cell. This feature of protein-partner exclusivity could be useful in *in vivo* applications, such as synthetic biology where a designed TF can work exclusively on an engineered target within a cell, thereby minimizing off-target effects,^62^ and in development of protein-based drugs that target a specific disease, without interactions with endogenous proteins that led to unwanted effects.^7^

We used rational design to generate mutants with the FosW LZ appended to Helix 2 of the **ME47/AA** bHLH domain. These mutants are termed “**MEF**” (<u>ME</u>47/AA-<u>F</u>osW). We examined the junction between the ME47 Helix 2 and FosW LZ by using crystal structures and the protein sequences to align leucines in heptad repeats at the dimerization interface. The LZ contains Leu every seventh amino acid. When two LZs dimerize, these heptad repeats align leucines at the hydrophobic protein-protein interface, thereby forming a parallel coiled coil.^63,64^ Charged residues often flank the hydrophobic core; a basic residue on one monomer can complement an acidic residue on its partner. This combination of Van der Waals and Coulombic interactions governs dimerization affinity and partner selection.

In MFW,^14^ swapping the native Max LZ for the FosW LZ was more straightforward to accomplish by aligning the Max LZ and FosW LZ sequences (Fig. S10B), but **ME47/AA** is a bHLH protein with no actual LZ element to swap with the FosW LZ. For **MEF**, the ME47 and Max-DNA crystal structures^13,21^ showed that the two protein structures overlay reasonably until V49 in Helix 2 of ME47 where they diverge (Fig. S12A). Hence, we chose V49 to be the C-terminus of ME47 Helix 2 to which we appended the FosW LZ; we used the same portion of the FosW LZ to generate MFW. We made **MEF** variants MEF+1 and MEF-1 to account for alignment of ME47 Helix 2 with FosW; because one turn of the α-helix is ∼3.5 amino acids, these three variants should essentially cover one turn (Fig. S12B). Testing the three variants should increase our chance of finding the optimal overall helical structure at the dimerization interface with aligned leucines, such that the hydrophobic interface extending from Helix 2 to FosW will not be out of register (Fig. S13).

In the B1H assay, **MEF** activated transcription from the E-box at a noticeably higher level than did **ME47/AA** (Fig. 4). Both MEF+1 and MEF-1 demonstrated modest activation from the E-box that was even weaker than **ME47/AA**’s activation from E-box, indicating that these mutants did not have the E47 HLH and FosW LZ joined together in the proper register to maintain the HLHZ dimerization interface. Using Western blot analysis, we observed that the amount of **MEF** protein expressed in the B1H assay surpassed that of **ME47/AA** (Fig. S14); this could arise from **MEF** being a more stable, better folded protein with slower degradation compared with **ME47/AA**.

**Figure 4.**
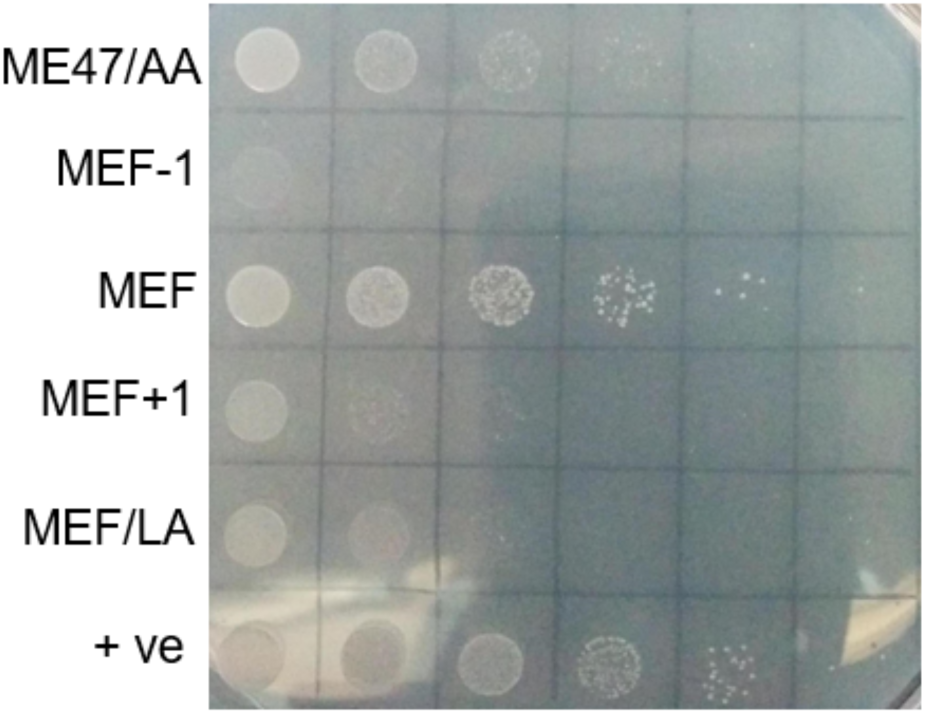
B1H assay comparing MEF variants to ME47/AA. Fusion of the FosW LZ out-of-register with the ME47 HLH (MEF-1 and MEF+1 variants) resulted in weaker transcriptional activation from E-box than that exhibited by **ME47/AA. MEF** produced the best signal observed in the B1H assay to date, validating the rationale for protein design. MEF/LA surprisingly showed activity greater than out-of-register MEF variants, despite having a LZ incapable of dimerization. Samples were plated on 5 mM 3-AT in 10-fold serial dilutions (10^−1^ to 10^−6^).

Consistent with activity measured in the cellular B1H assay, **MEF** exhibited excellent high-affinity, E-box-specific binding in quantitative EMSA (*K*_d_ 7.8 nM, 12-fold discrimination of binding to E-box vs. NS DNA, Table 1, Fig. 5). In comparison, **ME47/AA**’s binding to the E-box is 4-fold weaker with only 3-fold discrimination between the E-box and NS DNA targets, and MEF+1 and MEF-1 exhibit modest binding to the E-box that largely stems from nonspecific interaction with DNA (ratio of binding to E-box:NS DNA is 2:1). Moreover, Hill coefficient analysis of quantitative EMSA data demonstrated that **MEF** is clearly cooperatively bound to E-box as a dimer (Hill coefficient 2.3), whereas MEF+1, MEF-1 and **ME47/AA** all display noncooperative E-box-binding. Unusually, CD consistently showed markedly lower α-helicity displayed by **MEF** in comparison with **ME47/AA**, and we did not pursue CD for the MEF+1 and MEF-1 variants. However, the B1H and EMSA analyses confirm that both *in vivo* and *in vitro*, dimeric **MEF** binds to the E-box with high affinity and specificity.

**Figure 5.**
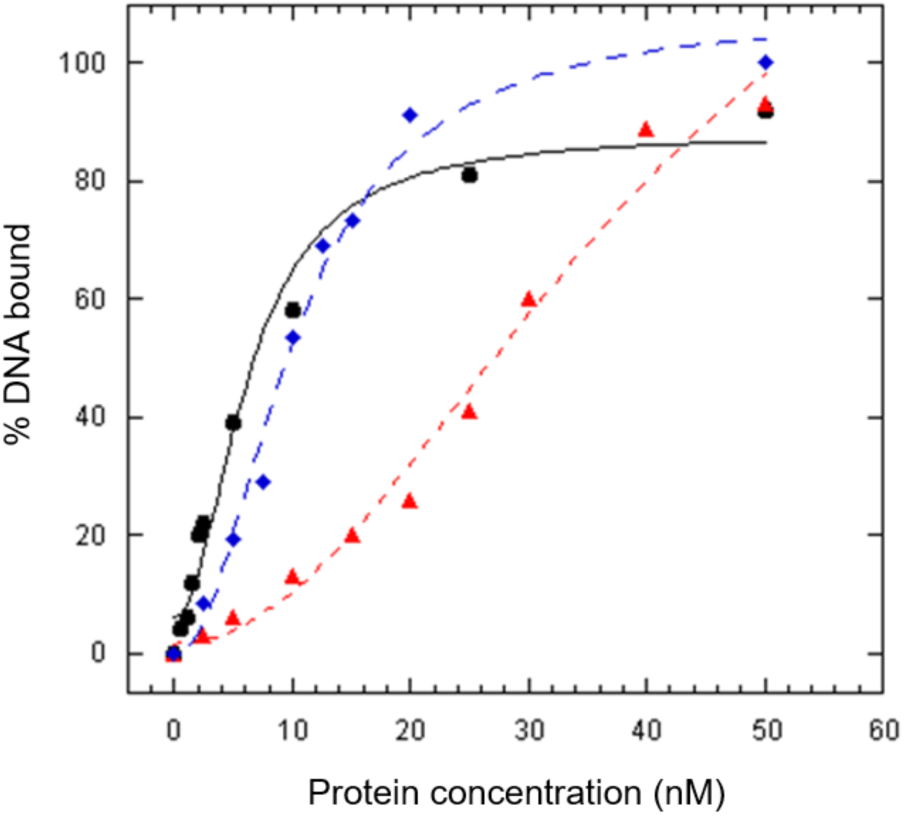
Representative equilibrium binding isotherms. Binding isotherms of E-box DNA bound to **MEF** (blue diamonds), MEF-1 (red triangles), MEF/LA (black circles). Each isotherm was obtained from an individual EMSA experiment. *K*_d_ values shown in Table 1 are averaged from individual isotherms. *R*^*2*^ values for all fits >0.99.

Given that Hill coefficient analysis indicated that **MEF** adopts dimeric, cooperative structure upon E-box-binding, we explored whether the FosW LZ was indeed responsible for homodimerization, as **MEF/AA**’s Hill coefficient is consistent with noncooperative E-box binding. To do so, we mutated the heptad Leu residues in the FosW LZ of **MEF** that form the hydrophobic interface that promotes coiled-coil dimerization; mutating these Leu residues to Ala minimizes their role in enhancing dimerization but maintains strong α-helical secondary structure, and we used this tactic previously on the FosW LZ in MFW.^14^ Interestingly, this Leu-to-Ala mutant MEF/LA showed reasonable transcriptional activation from the E-box in the B1H and a *K*_d_ value to the E-box approaching that of MEF (Fig. 5, *K*_d_ values 10.7 nM and 7.8 nM, respectively). MEF/LA is a more promiscuous DNA-binder, as shown by its relatively strong binding to NS DNA at *K*_d_ 44.5 nM, demonstrating that a significant component of MEF/LA’s binding to E-box is nonspecific. Hill coefficient analysis showed that MEF/LA is binding to the E-box as noncooperative monomers as expected given that the mutated FosW LZ no longer possesses Leu heptad repeats to promote the hydrophobic dimerization interface. CD analysis showed that MEF/LA and **MEF** possess the same levels of α-helical secondary structure, whether in the presence or absence of DNA (Table 1, Fig. 6); thus, the Leu-to-Ala mutations did not negatively impact protein structure and foldedness, just the ability to dimerize. Therefore, the FosW LZ in **MEF** is responsible for dimeric, cooperative binding to the E-box.

**Figure 6.**
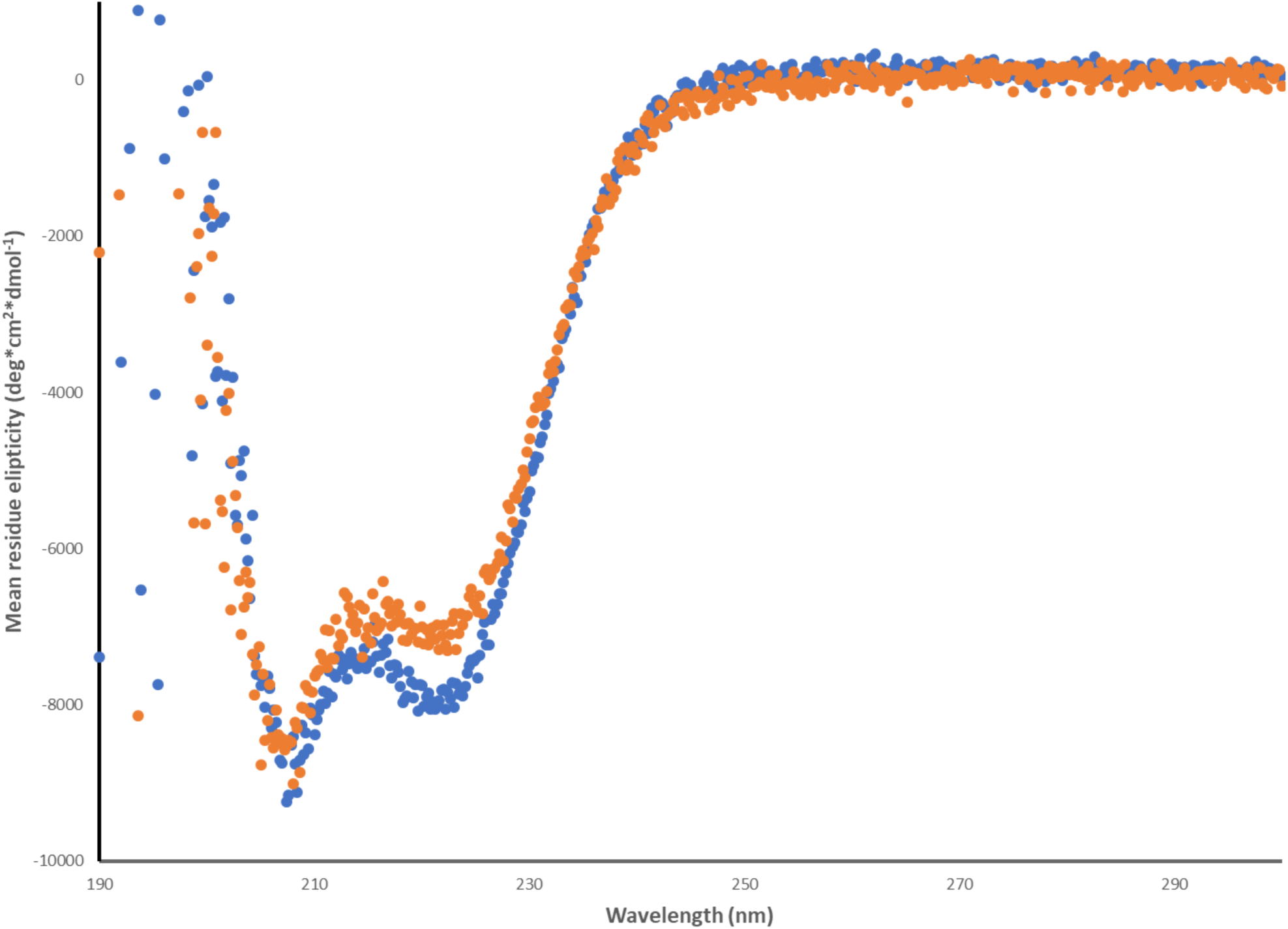
Circular dichroism spectra for MEF and MEF/LA. 2 µM **MEF** monomer (thus, 1 µM homodimer), orange circles; 2 µM MEF/LA, blue circles. Protein was resuspended in CD buffer (15 mM Na_2_HPO_4_, 5 mM KH_2_PO_4_, 50 mM NaCl, pH 7.4) and incubated for 1 hr, 37 °C. Each scan was carried out twice at 22 °C, scanning from 190 to 300 nm; scans were averaged and not subjected to smoothing. The buffer control was subtracted from each protein spectrum. Spectra for MEF with E-box or NS DNA are available in Supporting Information.

These data support that appending the FosW LZ to **ME47/AA** to generate **MEF** led to critical improved and desirable features: *1)* **MEF** binds to the E-box with *greater binding affinity* than does **ME47/AA** (*K*_d_ values 7.8 nM and 27.7 nM, respectively); *2)* **MEF** binds with *increased DNA sequence specificity* than does **ME47/AA** (E-box:NS DNA for **MEF** is 12:1 vs. 3:1 for **ME47/AA**); *3)* **MEF** binds to the E-box in dimeric, cooperative fashion, while **ME47/AA** does not; such cooperativity contributes to enhance **MEF**’s E-box-binding specificity and affinity. Most importantly, we find that **MEF** is a very stable and robust protein that is not prone to misfolding and aggregation, unlike ME47.

### The FosW LZ enhances both protein structure and DNA-binding function

We appended the FosW LZ to the **ME47/AA** bHLH to generate a bHLHZ structure for two main reasons: *1)* to enhance stability of the bHLH domain by nucleating α-helicity (structure), and *2)* to promote dimeric, cooperative binding to the E-box DNA target (function) toward compensation for mutation of Cys29 to Ala. Regarding structure, because ME47 can display instability, our logic was to append a LZ α-helix that can both promote and maintain protein helicity and foldedness. Previously in the case of bHLH/PAS protein Arnt, we replaced Arnt’s PAS module with the C/EBP LZ. The Arnt bHLH domain itself is capable of reasonably strong E-box binding *in vitro* (*K*_d_ 40 nM) but inactive *in vivo* in the yeast one-hybrid assay (Y1H), pointing to marked differences between the *in vitro* and *in vivo* assay data.^28^ However, fusion of the C/EBP LZ to Arnt bHLH restored E-box binding in the *in vivo* Y1H assay. In another designed protein, fusion of the c-Fos LZ to the Arnt bHLH domain also restored protein function in the Y1H.^14^ We surmised that in both of these cases, the appended LZ enhanced stable structure in the *in vivo* Y1H environment, but did not necessarily contribute to specific DNA-binding function, as fusion of the C/EBP LZ or the c-Fos LZ to the Arnt bHLH markedly weakened its binding to the E-box in quantitative fluorescence anisotropy titrations: these LZ fusions did not behave consistently in *in vivo* and *in vitro* assays. In contrast, MFW, with the truncated Max bHLH fused to FosW, demonstrated consistent activity *in vivo* and *in vitro*; MFW possesses good α-helicity as measured by CD and reasonable DNA-binding function both in the Y1H and quantitative fluorescence anisotropy.^14^

These previous observations indicate that LZs can serve as nucleation devices that encourage proper folding of the bHLH domain. Although **MEF** did not display increased α-helicity as measured by CD (Table 1, Fig. S15), we find that MEF has been extremely reliable to work with and produces reproducible results, similar to our workhorse protein MFW. Both **MEF** and MFW contain the FosW LZ.

The addition of the FosW LZ to **ME47/AA** also significantly enhanced binding to the E-box target. **MEF** displayed noticeably stronger binding to the E-box compared to **ME47/AA** in the B1H and four-fold enhanced binding by quantitative EMSA, with significantly increased specificity to the E-box vs. NS DNA. We surmise that **MEF**’s improved E-box binding also arises from the FosW LZ promoting dimeric structure capable of cooperative E-box binding. Although our original ME47 displayed dimeric structure by crystallography,^13^ we measured a Hill coefficient to E-box of 1.6, and all of the ME47 derivatives in Table 1 show largely noncooperative binding of two monomers to E-box. **MEF** is the only protein that conclusively shows cooperative binding to E-box.

### Evolution in nature and in the lab

We focus on transcription factors, because TFs are highly modular; for instance, just the DNA-binding domain or just the activation domain can be used and will still retain its native activity. In contrast, an enzyme’s active site comprises amino acids from various, distal regions of a protein’s sequence and cannot be sequentially linked into an “active-site domain.” Because our designs are franken-proteins—that is, modules from native TFs that are pasted together to generate an unnatural protein—they have not benefited from evolutionary pressure that can optimize structure and function. Moreover, these transcription-factor elements—originating from bHLH, bHLHZ, or bZIP— have been removed from their native structural context in which they evolved and then pasted into a new scaffold where they have not yet had a chance to co-evolve. For example, the Max basic region evolved in conjunction with the Max HLHZ, where both the HLH four-helix bundle and the LZ coiled-coil provide the dimerized structure that optimally positions the Max basic region in the DNA major groove to recognize the E-box sequence. However, the non-native combination of the Max basic region fused to the E47 HLH dimerization element, which is ME47, has not undergone the process of evolution, such as competing against other E-box-binding proteins and evolving under selective pressure. But PACE can provide a laboratory means for conducting Darwinian evolution.

Rational design did allow us to generate ME47, which is an effective inhibitor of breast cancer tumor growth in a mouse model and a somewhat tractable protein to work with in the lab.^43^ But we saw an opportunity to combine our rational approach with nonrational continuous evolution to improve ME47. PACE helped to fill the gap of finding non-obvious mutations that we would not have predicted or even tried in the lab: these mutations were captured under an evolutionary system providing constant mutagenesis and selective pressure for E-box binding function. In addition, our example above wherein the ME47 Cys29 may form a disulfide linkage, despite our crystal structure showing the two Cys29 thiols being too distant to react, demonstrates the protein flexibility and dynamism that can be difficult to anticipate by rational design.

Moreover, almost half of the residues in human TFs reside in intrinsically disordered regions,^65^ which can limit the utility of rational approaches in design of new proteins. These disordered regions are ubiquitously found in the eukaryotic proteome, and in particular, disordered structure in TFs can be beneficial to their DNA-binding functions: such versatile structures display flexibility, conformational adaptability, and exquisite ability to modulate on and off rates of DNA-binding necessary for fine-tuning gene regulation.^65,66^ The bHLH and bHLHZ TFs possess significant intrinsic order,^49^ including the HLH loop, which can critically contribute to a TF’s DNA-binding target affinity and specificity.^55^ Disordered protein structures have endured the rigorous process of natural selection and demonstrated their utility; for successful protein design, it is essential for us to exploit intrinsically disordered regions and proteins. Evolutionary methods that mimic the natural selection process, which has chosen intrinsically disordered regions/proteins time and again, are vital to successful protein design.

The *synergistic combination* of rational design and continuous evolution allowed us to truly improve ME47 to make **MEF**, which is markedly more robust and stable in our hands and possesses superior E-box binding function. Similar to our testing of ME47 as a cancer inhibitor,^43^ **MEF** could also serve as a protein inhibitor in human cells; as bHLH and bHLHZ proteins function as homo- and heterodimers in the cell, we were concerned that ME47, harboring the native E47 HLH domain responsible for E47 dimerization with protein partners, could dimerize with endogenous proteins in human cells. By designing franken-protein MEF whose dimerization and partner recognition depend on both the HLH and LZ, we rationally surmised that a dimerization interface comprising a mutated E47 HLH and non-native FosW LZ would likely avoid endogenous proteins and only homodimerize. Other systems for continuous evolution are being developed that should allow researchers opportunities and flexibility for incorporating evolution methods, including those for continuous evolution in human and mammalian cell systems.^67,68^ Such systems may facilitate development of proteins useful in human medical interventions, as well as veterinary care. As Max and E47 are found in humans and mammals, continuous evolution in these advanced eukaryotic systems may reveal mutations that allow MEF to perform better in mammalian/human cells, and can allow close examination of MEF’s protein-partner preferences under selective pressure in the presence of competing endogenous bHLH, bHLHZ, and bZIP proteins.

We have found that rational design and nonrational continuous evolution are complementary, synergistic methods toward our overall goal of protein design. As evolution methods are developing and taken up by more groups, we believe this two-pronged design strategy will show high utility and yield interesting and useful proteins, both scholarly and applied. Ultimately, a three-pronged approach that includes computational design will have the highest utility. Addition of computational approaches, most notably Rosetta (recently discussed in ref. 69), would make for a most powerful “trifecta.” As Kuhlman notes, natural proteins are apparently not optimally stable, and this lack of thermodynamic stability can be advantageous for proteins adopting diverse conformations and performing various functions.^69^ This concept is echoed by the wide natural abundance and utility of disordered proteins, on which evolution and computational approaches can have strong impact. Directed evolution and computational methods are two frontier fields that can greatly impact next-generation protein design.

## EXPERIMENTAL SECTION

### Protein nomenclature

Max bHLHZ refers to the human Max bHLHZ DNA-binding domain.^20,31^ E47 bHLH refers to the human E47 bHLH DNA-binding domain.^50^ FosW LZ was designed by Worrall and Mason.^15^ In FosW, 15 of the 35 amino acids in the native human c-Fos LZ were mutated to generate FosW, which homodimerizes; in contrast, c-Fos does not homodimerize, but heterodimerizes with bZIP partner c-Jun.^70^ Note that the original ME47 possesses Arg12 and Cys29 (Table 1, top line). ME47/RS refers to the ME47 mutant with Arg12 and Ser29, for example; thus, mutants are named “ME47/XX” and list the 12th and 29th amino acids after the backslash. (We note that ME47 could be termed “ME47/RC” in this nomenclature; however, as it is not a mutant, we simply refer to it as ME47.) For mutations at the 12th position, Arg12 is replaced by Cys, Ser, or Ala. For mutations at the 29th position, Cys29 is replaced by Ser or Ala. Table 1 shows all combinations of proteins with mutations at Arg12 and Cys29. We bolded the names of our main “take-home lesson” proteins **ME47/AA** and **MEF** to highlight them, because there are 18 unique proteins presented in this study. **ME47/AA** refers to ME47 with both Ala12 and Ala29 mutations (i.e., Arg12Ala, Cys29Ala). **MEF** is **ME47/AA** with the FosW LZ at the C-terminus. FosW was not appended to the C-terminus of **ME47/AA**, but rather the last seven amino acids of **ME47/AA** in Helix 2 were replaced by the FosW LZ. The FosW LZ comprises 35 amino acids,^15^ but we omitted its first four residues in our proteins.^14^ MEF+1 and MEF-1 are **MEF** variants that contain one extra or one less residue at the junction between ME47 and FosW, respectively; because the coiled coil dimer contains ∼3.5 amino acids/turn, **MEF** plus the two variants should cover one helical turn to enable alignment of the Leu residues in the FosW LZs for effective dimerization. MEF/LA is a control protein, in which the four Leu residues in the FosW LZ that enable dimerization are mutated to Ala (Fig. 1C). Although Ala is the best α-helix former and stabilizer,^57,58^ it does not promote LZ dimerization. All proteins are listed in Table 1 with B1H, CD, and *K*_d_ values. All protein sequences are shown in Figure 1C.

### Reagents

Reagents were purchased from BioShop Canada, enzymes were purchased from New England Biolabs (NEB), and oligonucleotides were synthesized by Eurofins Genomics or Integrated DNA Technologies unless otherwise noted.

### Bacterial strains

Bacterial strains were propagated in LB broth at 37 °C with shaking at 250 rpm. Ampicillin (50 µg/mL) and kanamycin (30 µg/mL) were added as appropriate. *Escherichia coli* DH5α (Stratagene) was used for standard cloning procedures. *E. coli* BL21(DE3)pLysS was used for the expression of His-tagged proteins. *E. coli* US0 cells (*ΔhisBΔpyrF*) were used to carry out the B1H assay.

### Plasmid construction

Detailed information on plasmid construction is provided in the Supporting Information. All new constructs were confirmed by dideoxynucleotide DNA sequencing performed at The Centre for Applied Genomics, The Hospital for Sick Children (Toronto, ON).

### Transformation, DNA, and plasmid preparation

Standard molecular cloning procedures were performed as described previously.^71,72^ Site-directed mutagenesis (SDM) was performed with Phusion high-fidelity DNA polymerase (New England Biolabs). SDM products and DNA fragments for cloning were purified using the Qiagen DNA purification and Qiagen gel purification kits (Qiagen, Mississauga, ON). For *E. coli*, plasmids were transformed by chemical transformation (TSS protocol^73^). Plasmids were purified using the QIAprep spin miniprep kit (Qiagen).

### PACE

The DNA-binding activity of ME47 was tested in the B1H system that originated in the Wolfe group^37,54^ before cloning into our PACE-B1H system. We cloned the sequence encoding ME47 into the pB1H2w2 plasmid C-terminal to the RNA polymerase omega-subunit and separated by a 21-residue linker; this construct is “pB1H2w2/ME47.” We cloned E-box constructs into the pH3U3 reporter vector 5’ to the -35 weak promoter of the *HIS3* reporter gene (Fig. S1). To optimize expression of the reporter gene by ME47, we tested a variety of E-box constructs, in which we altered *1)* the lengths of the nucleotide linkers between the E-box and - 35 weak promoter, and *2)* the flanking sequences outside of the core E-box CACGTG (discussed in Supporting Information, Fig. S2, Table S1).

To test the B1H signal of ME47, we co-transformed cells containing pB1H2w2/ME47 with the pH3U3/E-box reporter constructs and analyzed their B1H output using the *HIS3* growth assay (Fig. S3). We selected cells containing pairings of pB1H2w2/ME47 and pH3U3/E-box that showed maximum B1H signals and cloned their components into the respective PACE-B1H vectors, SP and AP, thereby generating SP/ME47 and AP/E-box. We tested these vectors in a plaque assay to assess plaque formation. The pair of SP and AP that together successfully formed plaques was used for PACE (Fig. S16).

The details of the PACE set-up are in preparation.^53^ Briefly, we sequentially transformed plasmids AP/E-box and MP6 (mutagenesis plasmid) into *E. coli* 1030. A single colony of AP- and MP6-transformed *E. coli* 1030 was used to prime the Chemostat containing ∼100 mL Davis Rich Media (US Biological) with appropriate antibiotics (ampicillin/chloramphenicol). Once the Chemostat was established, we connected and filled the lagoon with ∼30 mL *E. coli* culture. Media was continuously circulated through the Chemostat and Lagoon at 1 culture-volume/hr and 1.5 culture-volume/hr, respectively. We then primed the Lagoon with mature particles of SP-ME47 that are phage that were confirmed to produce plaques when allowed to infect permissive 1059 host cells. Arabinose was continuously added to the lagoon to maintain ∼10 mM arabinose concentration to induce mutagenesis. We ran the evolution experiment for 7 days after induction of mutagenesis. Every day, we collected and purified a sample of the Lagoon culture and tested in a plaque assay. Plaques were chosen to purify SP DNA for sequencing analysis (The Centre for Applied Genomics). Sequencing results were analyzed to identify any mutations that occurred in the DNA sequence encoding ME47 during PACE.

### Bacterial-one-hybrid assay (B1H assay)

The bacterial-one-hybrid system was adapted as previously described.^37^ The *E. coli* selection strain (US0 *ΔhisBΔpyrF*) was transformed with plasmids containing our reporter system in the pH3U3 vector, and the protein of interest was fused to the omega subunit of RNA polymerase in the pB1H2 vector. Optimization of the reporter system was carried out as described in the Supporting Information. The resulting transformants were plated on LB Amp+Kan plates, and overnight cultures were started from colonies on the plates. The overnight cultures were spun down and resuspended in NM-His**^+^** media (1× M9 salts, 40 mg/mL glucose, 10 µg/mL thiamine, 200 µM uracil, 200 µM adenine-HCl, 20 µM ZnSO_4_, 100 µM CaCl_2_, 1 mM MgSO_4_, <u>1% w/v Histidine</u>, 1× amino-acid mixture containing 17 of the 20 amino acids, excluding His, Met, and Cys),^37^ incubated for 1 hr, 37 °C, and washed in NM-His**^-^** media (same as the NM-His**^+^** media but *without* the addition of <u>1% w/v Histidine</u>) three times to remove any trace of histidine from the culture. The OD_600_ of the cultures were measured; the cultures were then diluted to a final OD_600_ of 0.1 and used to carry out serial dilutions from 10^−1^ to 10^−6^. 5 µL from each serial dilution was plated onto an LB plate that served as a positive control, and plated onto NM-His**^-^** plates (same as the NM-His**^+^** media but *without* the addition of His and adenine) that was supplemented with 3-amino-1,2,4-triazole (3-AT) in concentrations ranging from 1-10 mM to inhibit expression of the *HIS3* reporter gene. These plates were then incubated for 2 days, 37 °C to fully develop the colonies. The positive control consisted of US0 cells harboring pB1H2w2 containing Zif268, a zinc-finger TF, and pH3U3 containing the cognate sequence of Zif268 (GCGTGGGCG),^74^ while the negative control consisted of US0 cells harboring pB1H2w2 containing the DNA-binding protein under investigation and an empty pH3U3 vector.

### Cloning and generation of mutants

The DNA sequences encoding ME47 bHLH, MEF bHLHZ, and variants were ordered with *KpnI, XhoI*, and *XbaI* sites as shown in the Supporting Information. Cloning into the pB1H2 vector required digesting both the vector and duplex containing the protein with *KpnI* and *XbaI* in a sequential digest, with each digest carried out at 37 °C, 3 hrs after which the doubly digested pB1H2 was treated with Antarctic Phosphatase for 1 hr, 37 °C. The digested vector and duplex were then ligated using T4 DNA Ligase at 14 °C overnight. The same process was carried out for cloning into the pET28A(+) vector, with *KpnI* and *XhoI* used instead. The various mutants of ME47 and MEF were generated via site-directed mutagenesis. Primers and conditions used to carry out the mutagenesis are listed in the Supporting Information. Sequencing of the resulting cloning/SDM experiments was carried out by The Centre for Applied Genomics.

### Protein expression

The ME47 bHLH, MEF bHLHZ, and variants were bacterially expressed and purified following previously described methods with modifications described below.^5^

DNA fragments encoding the proteins in *E. coli* optimized codons were assembled and cloned into the pET28A(+) expression vector (Novagen; details of gene construction are given in the Supporting Information). These vectors were transformed into *E. coli* BL21(DE3)pLysS for protein expression. Typically, cells were grown in a 1 L culture where protein production was induced by adding 1 mM IPTG during the mid-log phase of growth (OD_600_ ∼0.6). After induction, the cells were harvested, sonicated, and purified using Co^2+^ metal affinity chromatography (TALON, Clontech) following the manufacturer’s protocol. The recommended buffers for TALON typically contain up to 5 mM β-mercaptoethanol (BME). However, 20 µL 1M DTT was used in lieu of BME due to problems previously encountered, where BME was found to be covalently linked to cysteine residues in our TFs.^29^ After TALON, proteins in the elution fraction were reduced by exposure to 10 mM DTT for 1 h, 37 °C and further purified by reversed-phase HPLC (Beckman System Gold) run on a semi-preparative reversed-phase C18 column (Vydac) with a gradient of acetonitrile and water plus 0.06% trifluoroacetic acid (v/v); the flow rate was 3 mL/min, and the gradient started at 0–20% acetonitrile over 20 min, followed by 20–60% acetonitrile over 45 min. Protein identities were confirmed by ESI-MS (Waters Micromass ZQ, Model MM1), and their concentrations were measured by UV/Vis spectrometry (Nanodrop 2000 Spectrophotometer, Thermo Scientific). Finally, the proteins were lyophilized in aliquots and stored at -80 °C. Immediately before EMSA, proteins were reconstituted to the desired stock concentration (typically 20 µM) in buffer and incubated for 1 h, 37 °C to maximize solubility.

### Electrophoretic mobility shift assay (EMSA)

Single-stranded oligonucleotides containing the desired protein-recognition site were synthesized with 6-carboxyfluorescein (6-FAM) incorporated at their 5’ ends (Fig. 1B). The 6-FAM-labeled oligonucleotides were annealed to their corresponding unlabeled complementary oligonucleotides by mixing the two in 10 mM Tris–HCl, pH 7, with the unlabeled oligonucleotide in 1.5x molar excess, heating at 95 °C, 10 min, and slow-cooling to room temperature. These annealed DNA targets were used in the following binding reactions with appropriate amounts of protein solution added to each reaction to cover the full titration range of the binding reaction. Protein-DNA binding reactions were performed in EMSA buffer (4.3 Na_2_HPO_4_, 1.4 mM KH_2_PO_4_, 150 mM NaCl, 2.7 mM KCl, 1.5 mM EDTA, pH 7.4, 2 mM DTT, 100 µg/mL BSA, 2 µg/mL poly dI–dC) and 2 nM 6-FAM-labeled duplex DNA in 30 µL total volume with 4 µL 30% ficoll added to the samples just before loading onto the gel. Prior to electrophoresis, samples were treated with the temperature-leap tactic (T-leap) to minimize protein misfolding and aggregation: protein solutions were incubated at 4 °C overnight followed by 30 min, 37 °C, and 1 hr, room temperature.^59,75^ The samples were loaded onto a pre-equilibrated native PAGE gel (10% poly-acrylamide, 0.5 % TBE), and run at 200 V for 5 min followed by 100 V for 25 min. The gels were visualized using the BioRad ChemiDoc MP Imaging System and analyzed, as detailed previously.^29,76^ A brief summary follows. Imagelab software (Version 5.2) was used for densitometric analysis for calculation of the bound DNA fraction (*θ*_app_) for each lane, where the bound DNA fraction is the intensity of the band corresponding to the protein-bound DNA divided by the sum of the intensities of the bands corresponding to the protein-bound DNA and free DNA. The bound DNA fractions were fit to the equation below using KaleidaGraph software (Version 4.5) to calculate the apparent *K*_d_ value, as described previously.^76^

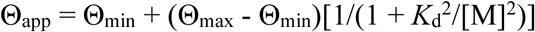

All data sets fit to Eq. 1 had *R*^*2*^ values >0.99. Each *K*_d_ value was determined from the average of two independent experiments. Representative EMSA are shown in Fig. S11.

### Circular dichroism (CD)

Single-stranded oligonucleotides containing the desired protein-recognition site were annealed by mixing the complementary oligonucleotides in 10 mM Tris– HCl, pH 7, heating at 95 °C, 10 min, and slow-cooling to room temperature. The protein was resuspended in CD buffer (15 mM Na_2_HPO_4_, 5 mM KH_2_PO_4_, 50 mM NaCl, pH 7.4) to a final stock concentration of 20 µM and incubated for 1 hr at 37 °C. The protein was then diluted to 2 µM final concentration in 2 mL final volume with CD buffer (and DNA duplex if applicable, which was diluted to 2 µM final concentration). The protein-DNA mixture was treated with the T-leap, as described above. CD was performed on an Aviv 215 spectrometer with a suprasil, 10 mm path-length cell (Hellma) at 22 °C. Spectra were acquired between 180 and 300 nm at 0.2 nm increments and a sampling time of 0.2 s. Each spectrum was the average of two scans with the buffer control spectrum subtracted. Data obtained were not smoothed. Protein α-helix content was calculated by the method of Chau and coworkers.^77^ Briefly, percent α-helix content was determined by assuming only α-helical content (i.e., assuming no β-sheet structure) and using the equation:

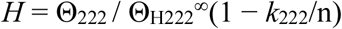

where *H* is the percent helicity, Θ_222_ is the mean residue ellipticity at 222 nm, Θ_H222_∞ is the reference value for a helix of infinite length, *k*_222_ is a wavelength-dependant constant, and n is the number of amino acids in the protein.

## Supporting information

Info about PACE, B1H, EMSA, CD, Kd values

## Acknowledgement

We appreciate assistance from Duan Fang Tan, Sama Abdulnabi, Esther Jung, and Mercy Daka. We are grateful to CHRP (Collaborative Health Research Project), NSERC Discovery Grant, and the UTM VP Research and UofT XSeed for support.

## Supporting Information

PACE details. Additional DNA target sequences and data. Primer sequences and cloning details. Representative B1H, CD, EMSA data.

## Notes

The authors declare no competing financial interest.

