## Supplementary material for "Combining rational design and continuous evolution on minimalist proteins that target DNA": Info about PACE, B1H, EMSA, CD, Kd values

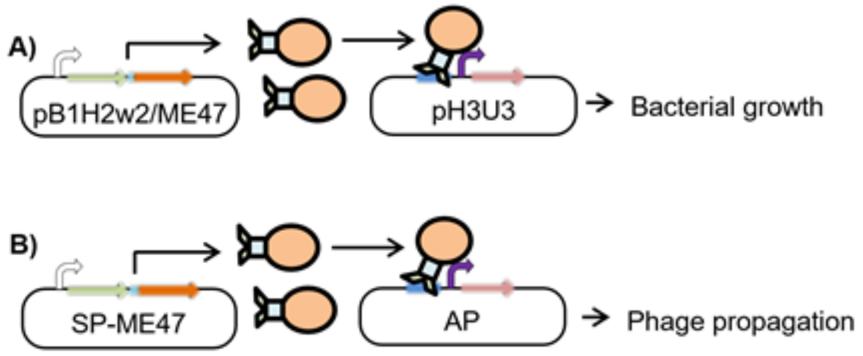


**Figure S1. Schematic diagram illustrating A) B1H system and B) PACE-B1H system**. The omega subunit of RNAP (RNA Polymerase) fused to ME47 (green and orange, respectively) acts as an activator domain (AD). ME47 binds to the E-box (blue) present on the pH3U3/AP vector (AP, accessory plasmid) and positions the omega subunit so that it can recognize the -10/-35 elements of the weak lac promoter (purple). This leads to expression of the downstream gene (beige) that is needed for *E. coli* growth and phage propagation. (Adapted from Popa, Thuronyi, Inamoto, & Shin, manuscript in preparation.)

```
-7 AT E-box    GCGGCCGCTGCAGGAACCACGTGGTGGAATTCTTTACA
-7 GC E-box    GCGGCCGCTGCAGGAGCCACGTGCGGGAATTCTTTACA
```

**Figure S2. Reporter system used for B1H/PACE experiments.** The reporter system consists of an E-box (yellow) positioned upstream of the -35 element of the weak lac promoter (green) that is separated by a spacer of a predefined length (underlined; we count from the beginning of the -35 element to 2 bp outside of the core E-box). The sequences flanking E-box can play a role in a transcription factor's (TF) DNA-binding affinity and specificity. In particular, we have found that nucleotides specifically 2 bp outside of the core E-box (red) affect protein-DNA recognition. We used spacer length and the red bp in naming the reporter system. For example, the top sequence is the "-7 AT E-box," where "-7" refers to the number of nucleotides between the base highlighted in red at 3' end of the E-box and the -35 element (i.e., the spacer), and "AT" refers to the nucleotides (red) that we have found to impact protein binding affinity and specificity. Thus, we experimented with both A/T and G/C bp positioned 2 bp from the E-box.

The weak lac promoter regulates the expression of downstream *HIS3* and gIII reporter genes that form the basis of the reporter systems for the B1H and PACE, respectively. The length of the spacer is such that it will allow the TF-RNAP fusion protein to bind the E-box and interact with the promoter for optimal expression of the reporter. Depending on the size of the fusion protein, different spacer lengths can be tested to optimize the system. *For this paper, all experiments done with the* ***E-box*** *refer to the* ***-7 GC E-box*** *shown here and in Figure 1B*.

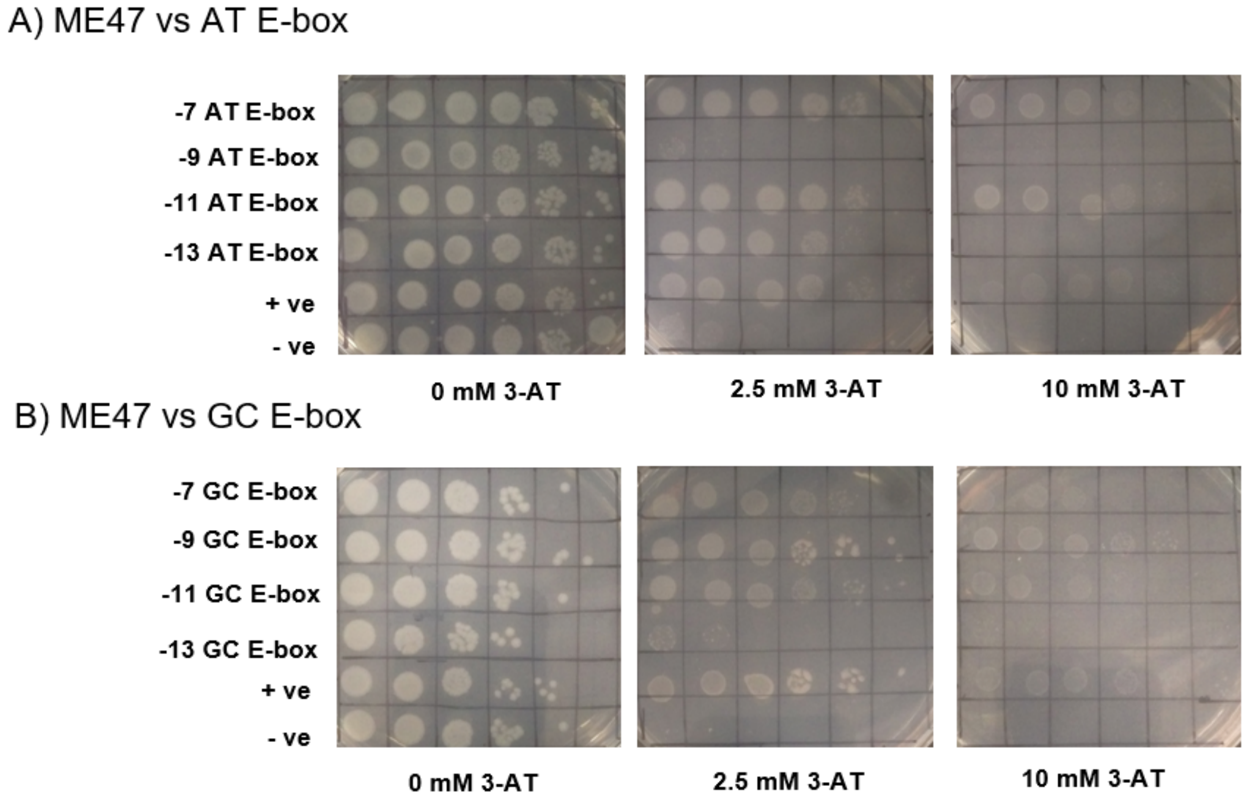


**Figure S3. Representative B1H assays for the GC and AT E-box libraries.** The libraries spanned 7 to 25 nucleotide spacers increasing in increments of 2 nucleotides (shown here are 7, 9, 11, 13 nucleotide spacers for both AT and GC E-boxes). The ideal E-box reporter would give a signal similar to the positive control even at higher concentrations of 3-AT, unlike the -9 and -13 AT E-box constructs shown in the 2nd and 4th rows of the top panel, as an example. We plated serial 10-fold dilutions from left to right ($10^{-1}$ to $10^{-6}$). The positive control consisted of US0 cells transformed with Zif268 and its cognate DNA in lieu of the E-box, while the negative control consisted of US0 cells transformed with Zif268 and empty pH3U3.[1] Omega-ME47 produced strong signals from the -7 and -11 AT E-box constructs and -7, -9 and -11 GC E-box constructs, making these promising candidates for the PACE-B1H. (Adapted from Popa, Thuronyi, Inamoto, & Shin, manuscript in preparation.)

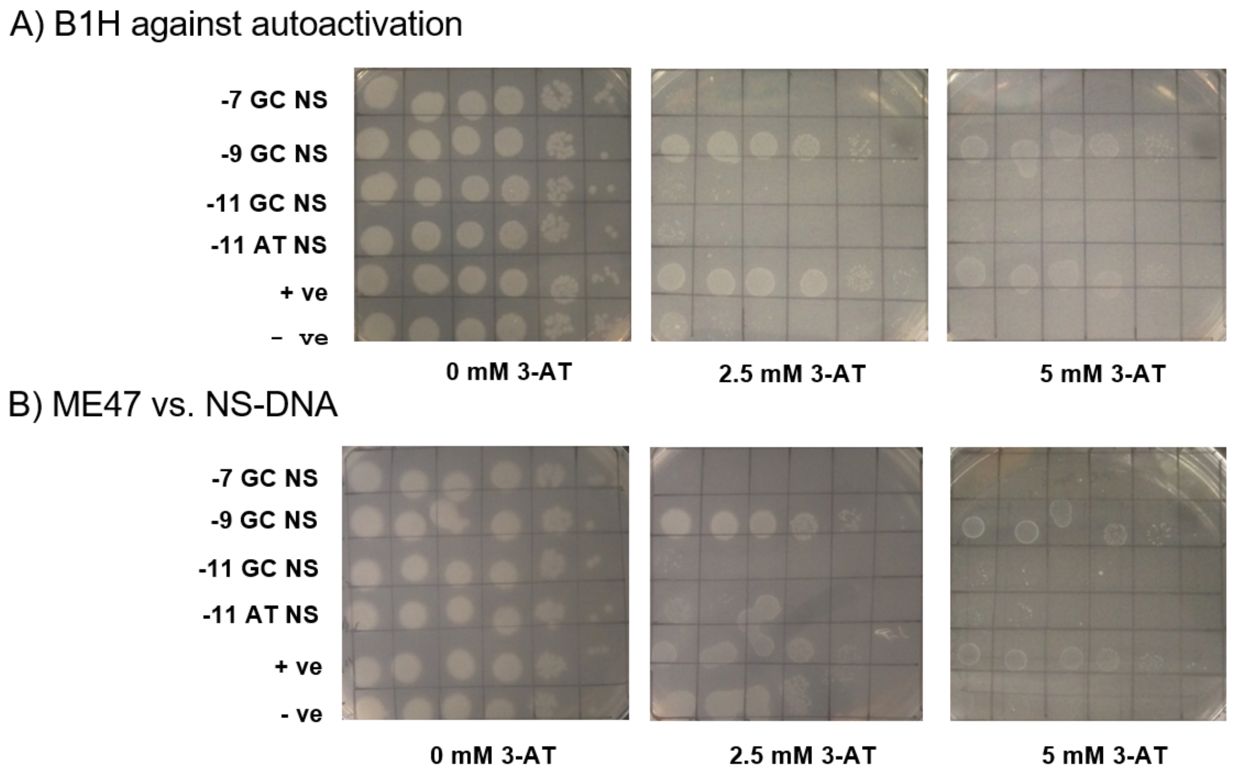


**Figure S4. A. B1H assays to test autoactivation.** The autoactivation B1H consisted of US0 cells transformed with the various library members on the pH3U3 plasmid *without* the ME47-omega subunit fusion on the pB1H2w2 plasmid (shown here are library members with -7 GC, -9 GC, -11GC, and -11AT nucleotide spacers for both conditions). In the absence of the ME47-omega subunit fusion protein, no signal should be seen. As such, the -9 GC E-box in the 2nd row would be considered an autoactivating E-box, while the -7, -11, and -13 GC E-boxes in the top row are not autoactivating. **B. B1H assays to test nonspecific DNA binding.** The nonspecific B1H consisted of US0 cells transformed with the ME47-omega subunit fusion on the pB1H2w2 plasmid and the same spacer library members used in Fig. S3A on the pH3U3 vector whose E-box was replaced with NS DNA (5'-TCCAAG in lieu of 5'-CACGTG). The controls were the same as described earlier. The -7 GC, -11 AT and -11 GC constructs (1st row, 3rd row, 4th row) did not produce a signal in the nonspecific B1H, making them promising candidates for use in the PACE selection circuit. (Adapted from Popa, Thuronyi, Inamoto, & Shin, manuscript in preparation.)

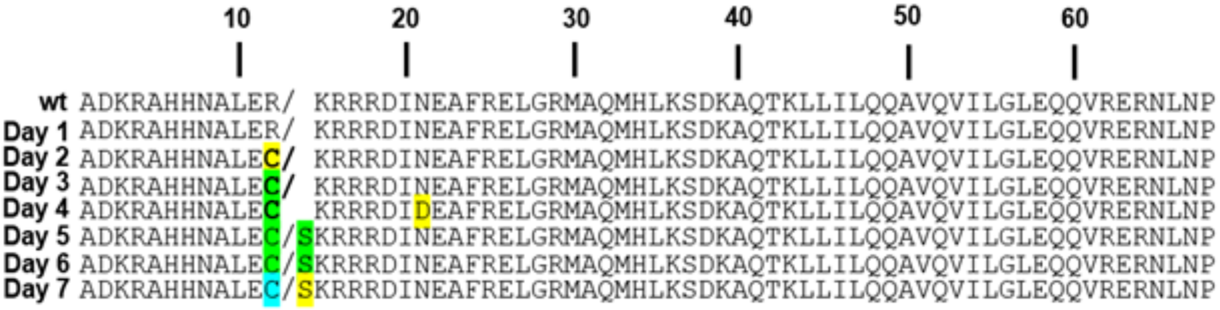


**Figure S5. Representative sequences of ME47 variants throughout the PACE run.** Mutations highlighted in yellow were present in <50% of the sampled population for that day. Mutations shown in green were present in 50% of the day's population, while mutations in blue were present in >75% of the day's population. The Arg12Cys and Arg12Ser mutations in ME47 dominated the Lagoon by day 6 of the PACE run. The Asn22Asp mutation did not persist in the Lagoon and was not explored, as we presumed it to be a chance mutation with no impact on phage propagation. For each day, 5 clones were sequenced.

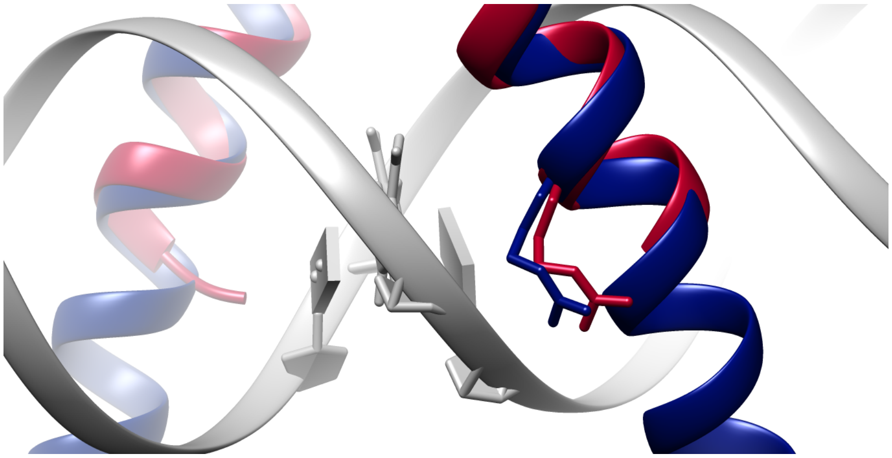

**Figure S6. Comparison of Max and ME47 basic regions.** Arg12 in the Max crystal structure (navy, PDB: 5EYO[2]) makes a DNA phosphodiester contact with the central bolded G in the E-box CAC**G**TG, which is why the mutants that arose in PACE at this position surprised us.[3] Our ME47 crystal structure is shown in red (PDB: 3U5V[4]). The loss of this contact should have resulted in decreased affinity and possibly specificity; we did not expect it to be selected for in PACE, but the opposite happened. Previously, we showed that mutating 24 of 27 Arg residues to Ala in the basic region of a bZIP protein resulted in proteins that retained native bZIP structure and AP-1 DNA-binding function.[5] We reasoned that a similar outcome occurred here. Alternatively, mutating the Arg12 position could favorably alter the junction between the Max basic region and the E47 HLH; Arg12 is only 3 bp from the junction with the E47 HLH. Chimera v. 1.11.2.[6]

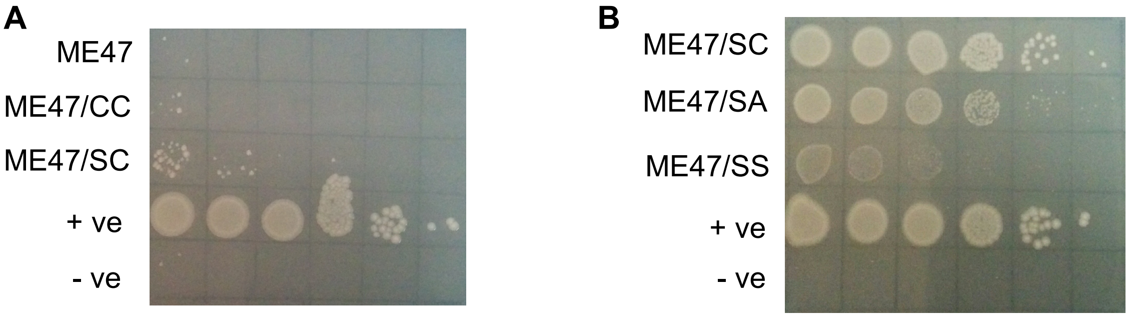


**Figure S7. A. B1H testing ME47/SC and ME47/CC mutants binding to NS DNA.** ME47/SC displayed slight nonspecific activity (3rd row), while ME47 and ME47/CC showed no nonspecific activity. These results are representative for all variants of ME47 and **MEF** (i.e., little or no binding to NS DNA), which suggests that the signal observed from our E-box containing reporters are the result of a true TF interaction with the E-box. Samples were plated on a 2.5 mM 3-AT plate, NS DNA cloned into pH3U3 reporter was cagt**tccaag**gc, with the sequence in bold replacing the E-box in the reporter construct. **B. B1H assay on Cys29 mutants of ME47/SC.** To eliminate unwanted Cys thiol side-chain reactivity, we rationally replaced Cys29 with Ser or Ala in mutant ME47/SC (Arg12 mutated to Ser). The Ala29 mutant ME47/SA gives a noticeably decreased signal relative to ME47/SC, while the Ser29 mutant ME47/SS gives surprisingly poor signal. *Ser29 mutants consistently showed poor activity in the B1H and were not further studied.* In PACE, ME47's Arg12 mutated to either Ser or Cys (we did not pursue Cys due to side chain reactivity). Both ME47/SC and ME47/SA show that mutation of Arg12 to Ser gives ME47 mutants that are highly active in the B1H assay. The six columns from left to right show cell growth in 10-fold serial dilutions ($10^{-1}$ to $10^{-6}$) on 20 mM 3-AT.

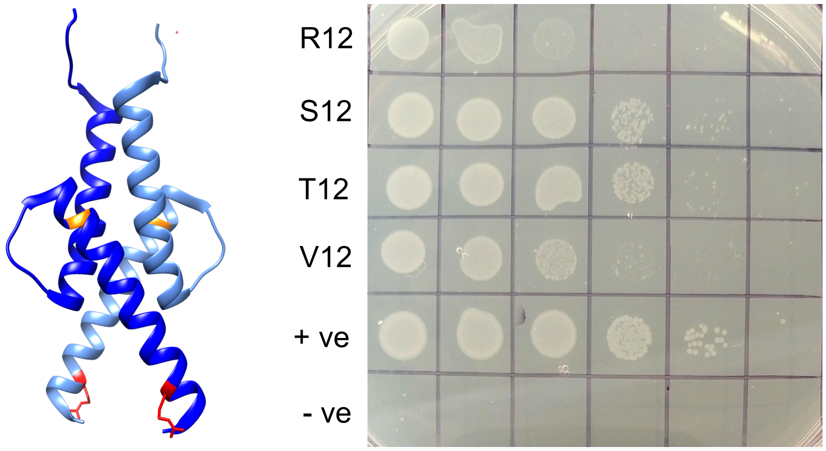


**Figure S8. Effects of rationally designed mutations at position 12 of ME47. Left.** ME47 (PDB: 3U5V[4]) shown as a homodimer (light and dark blue ME47 monomers) with Arg12 shown in red and Cys29 in orange. **Right.** To investigate the nature of the Arg12 mutation, Thr and Val were tested at position 12. ME47 Arg12Thr behaved similarly to ME47 Arg12Ser, which did not surprise us considering the structural similarity of the side chains. ME47 Arg12Val gave a weaker signal but was still stronger than the original ME47. A smaller residue at position 12 is essential for the observed gain in function, but a hydroxyl/sulfhydryl group also appears to be important. Our working theory is that a smaller residue at position 12 that is near the junction between the Max basic region and E47 HLH improves protein stability.

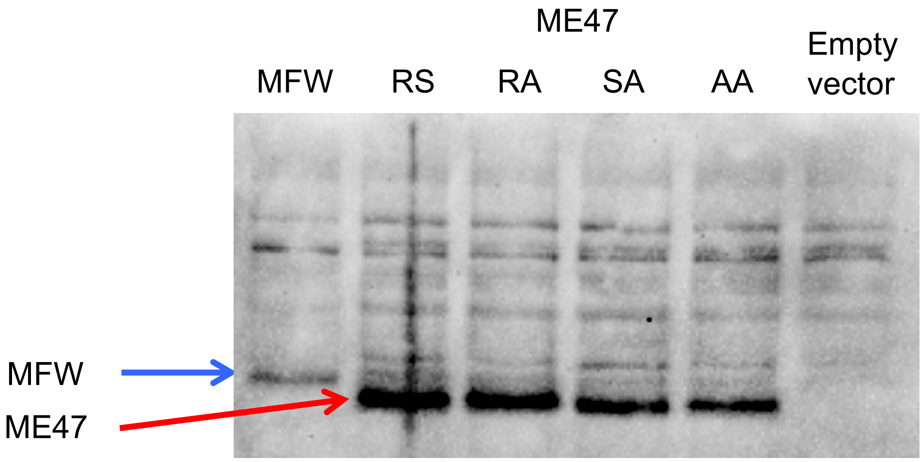


**Figure S9. Western blot analysis of MFW and ME47 expression levels.** US0 cells used for the B1H were transformed with plasmid, protein expression was induced with IPTG, and cells were lysed after a 2 hr induction period. Primary antibody was rabbit anti-ME47 polyclonal specific to the N-terminal Max basic region; secondary antibody was goat anti-rabbit HRP-conjugate.

From the overnight blot, the amount of ME47 and variants present in the cells exceeds that of MFW. Given these results, we were even more surprised that MFW outcompeted ME47 and its variants in the competition PACE. Not only does MFW have ~15-fold lower binding affinity to E-box than does ME47, but it is also much less abundant in the cell.

MFW overexpression is not the explanation for why MFW dominated the competition PACE. This suggests that MFW's "fitness" is due to some other beneficial feature it possesses over ME47. We hypothesize that MFW is more stable, robust, and well-folded. MFW always performs reliably in the lab, and it is very tractable and consistent.

At the conclusion of the competition PACE (3 days of PACE), 10 plaques were sequenced. All 10 plaques derived from cells infected with SP-MFW.

**A**

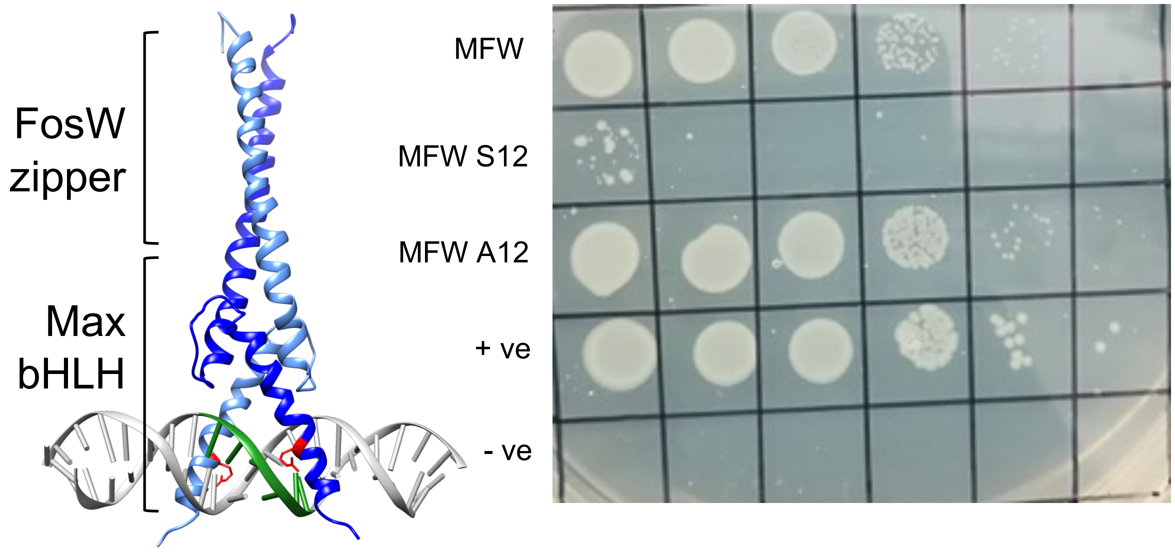


**B**

Max: -AQILDKATEYIQYM-RRKNDT**H**QQDIDD**L**KRQNAL**L**EQQVRA**L**EKARSSAQLQT

FosW: AASLDEL-QAEIEQ**L**EERNYA**L**RKEIED**L**QKQLEK**L**GAP

MFW: -AQILDKATEYIQYM-QAEIEQ**L**EERNYA**L**RKEIED**L**QKQLEK**L**GAP

**Figure S10. A. PACE-derived mutations are *not* interchangeable between similarly related proteins.** Given that MFW is also a "franken-protein" like ME47 and that it also contains the Max basic region (MFW = Max bHLH + FosW LZ),[7] we explored whether mutations at MFW's Arg12 would yield results similar to those for the ME47 mutants at Arg12. MFW with Arg12Ser has virtually no activity unlike the same mutation in ME47 that was uncovered by PACE, while MFW with Arg12Ala has activity comparable or better than original MFW, similar to the same mutation in ME47. Interestingly, Arg12 in MFW can be mutated and still maintain activity, suggesting that this Arg12-DNA phosphodiester interaction is not essential. The MFW model was generated using I-TASSER.[8] **B. Creation of MFW.** We aligned the Max bHLHZ and FosW LZ to identify the optimal position to make the LZ swap. In the top sequence, Max's Helix 2 sequence is underlined, and the dash separates Helix 2 from the LZ. In MFW, the full Max Helix 2 is preserved, while the first 7 aa of FosW were omitted. Residues in bold correspond to Leu residues (Max has one His) that comprise the heptad repeats for LZ coiled coil dimerization.

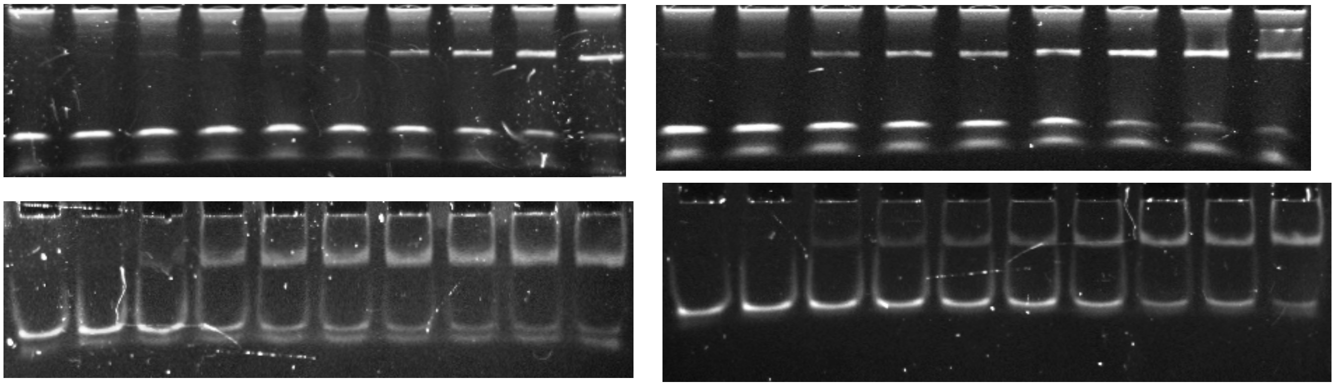

**Figure S11. Representative EMSA for ME47/AA and MEF proteins.** The $K_d$ value for each EMSA is given below in parentheses. The $K_d$ values in the Tables are the averages of $K_d$ values from two independent EMSA experiments.

***Top left***: **MEF** bound to GC E-box ($K_d$ = 7 nM); monomeric protein concentrations from left to right are 0, 0.5, 1, 1.5, 2, 2.5, 5, 10, 25, 50 nM.

***Top right***: **MEF** bound to GC NS ($K_d$ = 97 nM); monomeric protein concentrations from left to right are 25, 50, 60, 70, 80, 90, 100, 150, 200 nM.

***Bottom left***: **ME47/AA** bound to GC E-box ($K_d$ = 29 nM); monomeric protein concentrations from left to right are 5, 10, 15, 18, 21, 24, 27, 30, 33, 40 nM.

***Bottom right***: **ME47/AA** bound to GC NS ($K_d$ = 104 nM); monomeric protein concentrations from left to right are 20, 40, 50, 60, 70, 80, 100, 150, 200, 250 nM.

**Discussion of cooperativity & Hill coefficients.** E-box has two half-sites: each half-site can be bound by one protein monomer. With Hill coefficient ~1, binding to DNA is *noncooperative*, i.e., protein binding to one half-site does not affect the other half-site.[9] When the Hill coefficient is >1, this is positive cooperativity, which we observe in ME47, ME47/SC, and MEF with Hill coefficients 1.6, 1.5, and 2.3, respectively. For systems showing positive cooperativity, the Hill coefficient informs on the lower limit of the number of interacting sites.[10] Thus, these Hill coefficients of 1.5-2.3 indicate ~2 cooperatively interacting sites, consistent with the E-box comprising two abutting half-sites. The rest of our proteins display noncooperativity (Hill coefficients ~1).

We recognize that Hill coefficients can give a range of values, and that it is not entirely reliable, as discussed by Cattoni *et al.*[10] For instance, in the classic case of oxygen binding to hemoglobin,

Hill coefficients range from 2.8-3.4, depending on conditions, type of experiment, etc.

If our proteins were binding as one monomer on one E-box half-site followed by another monomer uncooperatively binding on the other E-box half-site, we would see this on the gels. We would observe one band for unbound DNA, another band for monomer-bound DNA, followed by a third band growing in for two monomers bound to DNA. But we only observe all-or-nothing binding, i.e., unbound DNA or two monomers/one dimer bound to DNA. All of our proteins show only two bands in EMSA, so we know that two monomers/one dimer are bound in the second, higher band.

This is consistent with our work on bZIP derivatives of GCN4,[11] and others' kinetics studies on bZIP proteins Jun and Fos, and bHLHZ proteins Max, Mad, and Myc,[12-16] showing that bZIP and bHLHZ proteins bind to DNA by a cooperative *monomer pathway* where one monomer binds to DNA and cooperatively, rapidly assists the second monomer onto DNA. Because the second monomer goes onto DNA so rapidly, effectively what is observed is dimerically bound DNA, i.e., two-state binding with unbound DNA and dimer-bound DNA. No monomer-bound DNA is observed. (However, Lavigne's lab showed evidence that with Max, Mad, and Myc, protein *heterodimers* formed first and then bound to DNA, so this is a *dimer pathway*.[17]) Both the dimer and monomer pathways give the same thermodynamic result (dimer bound to DNA target) and are energetically equivalent, while kinetics dictates preference for the monomer pathway.[16]

For our proteins with Hill coefficients ~1 (noncooperative binding), we do not have the actual monomer pathway, which is still *cooperative*. We are seeing two monomers without cooperativity bound at the E-box, so thermodynamically, this looks like a dimer on EMSA. Only MEF gives Hill coefficient ~2, indicating dimeric cooperativity when binding to E-box; ME47 and ME47/SC exhibit positive Hill coefficients ~1.5 indicating some contribution from cooperativity when binding to DNA.

**A**

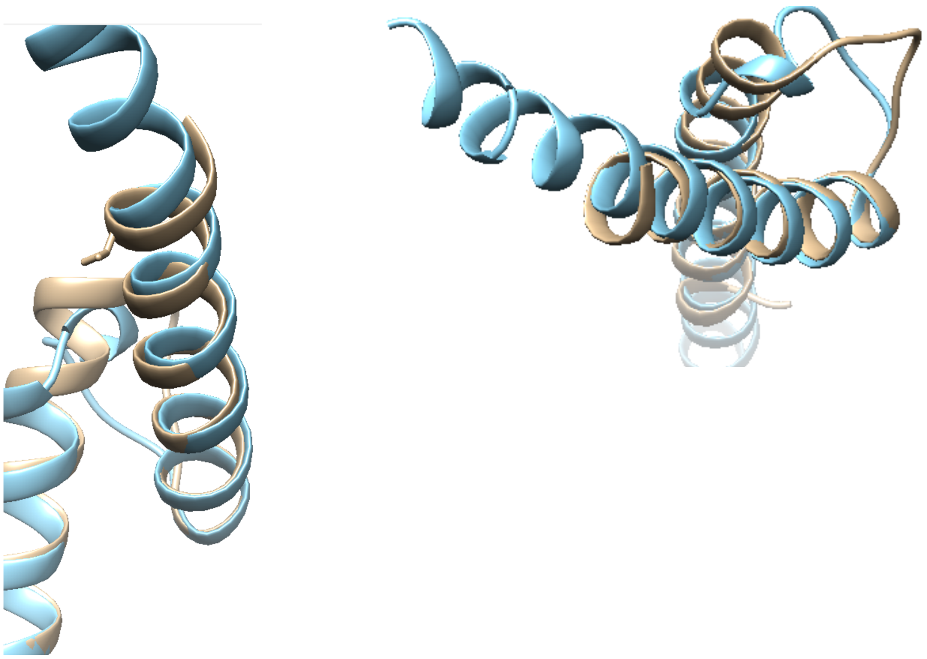

**B**

| | |
|---|---|
| ME47/AA | ADKRAHHNALEAKRRRDINEAFRELGRMAQMHLKSDKAQTKLLILQQAVQVILGLEQQ**V**RERNLNP------------------------ |
| MEF +1 | ADKRAHHNALEAKRRRDINEAFRELGRMAQMHLKSDKAQTKLLILQQAVQVILGLEQQ**V**QAEIEQLEERNYALRKEIEDLQKQLEKLGAPLE |
| MEF | ADKRAHHNALEAKRRRDINEAFRELGRMAQMHLKSDKAQTKLLILQQAVQVILGLEQQ**V**-AEIEQLEERNYALRKEIEDLQKQLEKLGAPLE |
| MEF/LA | ADKRAHHNALEAKRRRDINEAFRELGRMAQMHLKSDKAQTKLLILQQAVQVILGLEQQ**V**-AEIEQ**A**EERNYA**A**RKEIED**A**QKQLEK**A**GAPLE |
| MEF -1 | ADKRAHHNALEAKRRRDINEAFRELGRMAQMHLKSDKAQTKLLILQQAVQVILGLEQQ--AEIEQLEERNYALRKEIEDLQKQLEKLGAPLE |

**Figure S12. A. Rational design to create MEF.** Aligned crystal structures of ME47 (tan, PDB: 3U5V[4] and Max (blue, PDB: 1HLO[18]) visualize where to attach the FosW LZ. The mismatch in Helix 2 between the two proteins becomes noticeable at Val49 (Val side chain shown on left); we decided to fuse the FosW LZ around Val49. Because this was a rough approximation of the Helix 2/LZ junction, we created variants of the junction in order to align ME47 Helix 2 and FosW LZ in proper register for optimal dimerization. **B. Sequences of the MEF variants**. The residue shown in bold is Val49, where Helix 2 of ME47 was truncated and the FosW LZ was fused, as for MFW in Figure S9. **MEF** showed optimal DNA-binding function in the B1H and EMSA. MEF+1 and MEF-1 variants showed weaker function that we attributed to the ME47 HLH and FosW LZ being out of register. Chimera v. 1.11.2.[6]

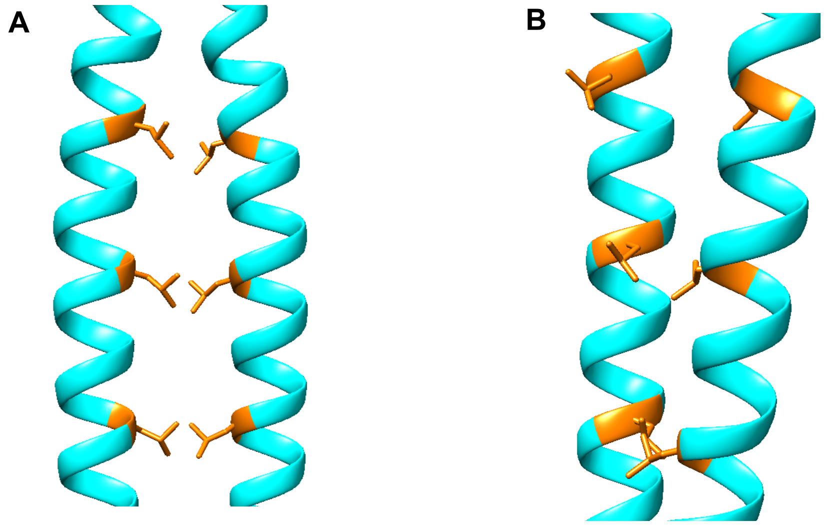


**Figure S13. Visualization of in-register and out-of-register LZ coiled coils. A.** When the LZs are in register, the hydrophobic leucines (orange) interact at the coiled coil interface to permit dimerization. **B.** When the LZs are out of register via addition/removal of residues from the LZ, there is no hydrophobic coiled coil interface, which impedes dimerization. Chimera v. 1.11.2.[6]

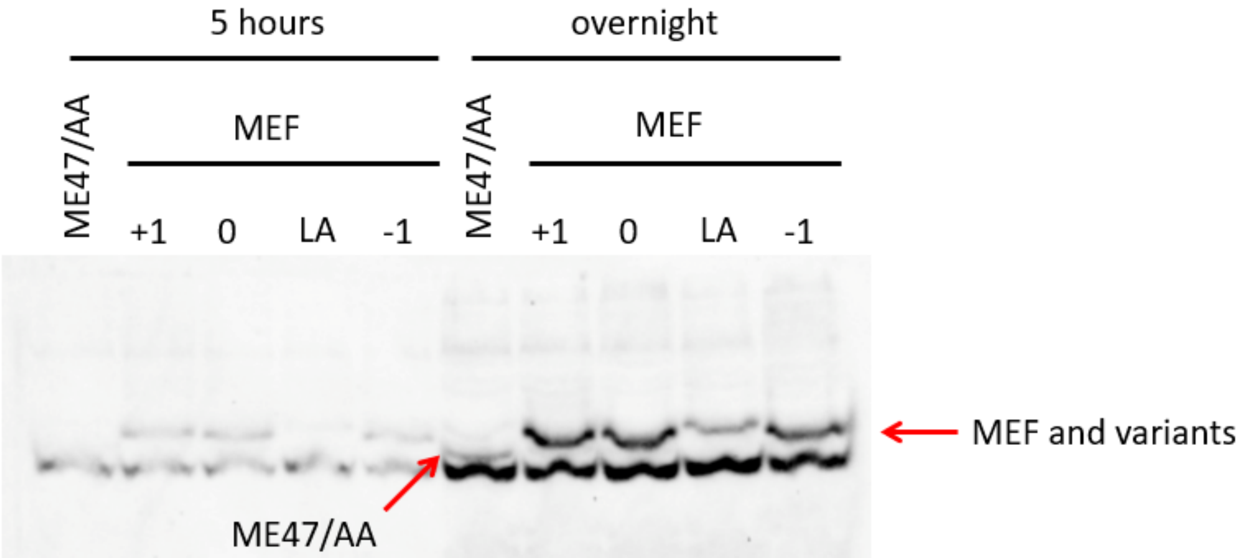


**Figure S14. Western blot testing ME47/AA and MEF expression levels.** US0 cells used for the B1H were transformed with plasmid, protein expression was induced with IPTG, and cells were lysed after a 5 hr or overnight induction. From the overnight blot, the amount of **MEF** and variants present in the cells dwarfs that of **ME47/AA**, which may result from **MEF** and variants being more stable, folded, and less susceptible to degradation than **ME47/AA**, which likely represents all ME47 variants regarding stability. Primary antibody was rabbit anti-ME47 polyclonal specific to the N-terminal Max basic region; secondary antibody was goat anti-rabbit HRP-conjugate. **MEF** indicated as "0," MEF+1 and MEF-1 as "+1" and "-1," and MEF/LA as "LA."

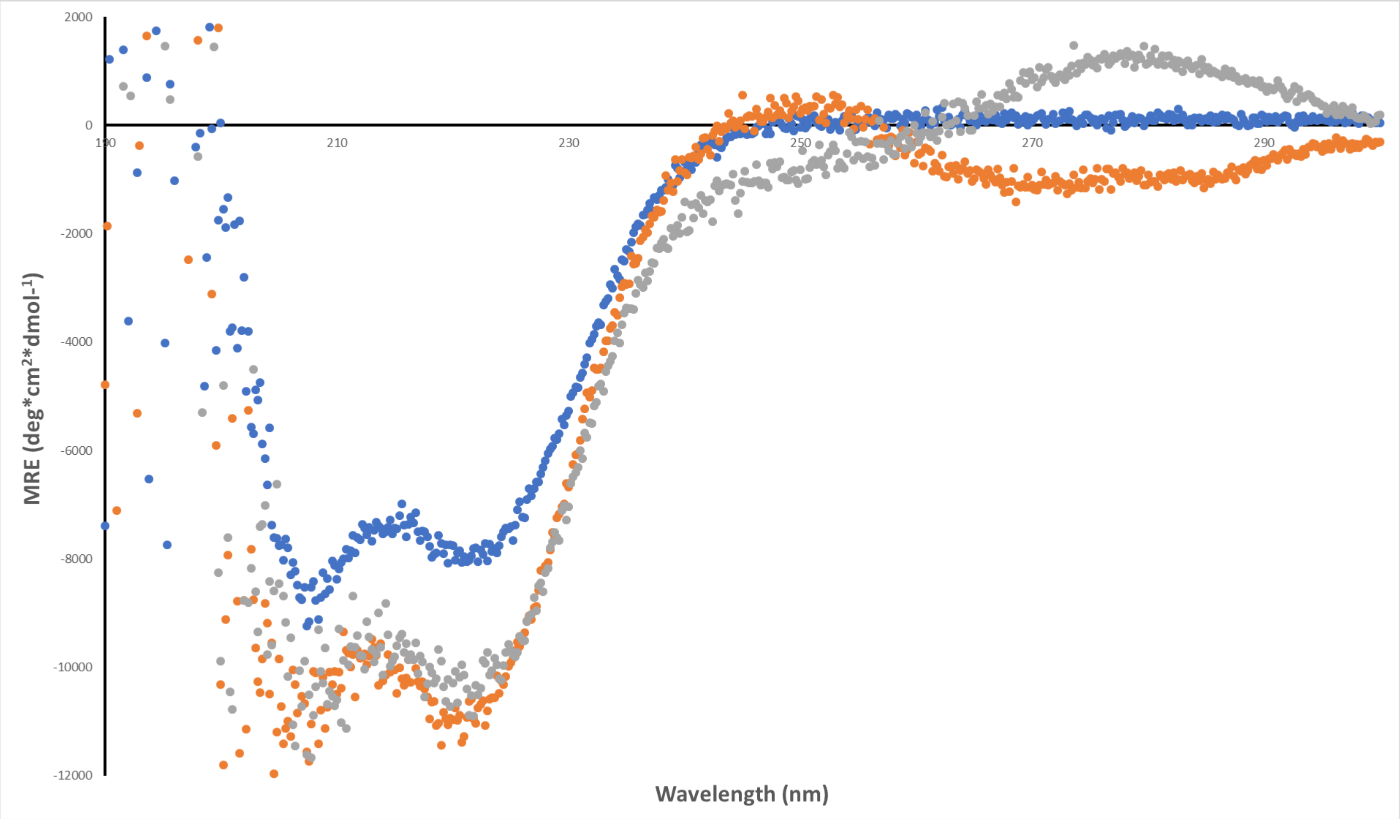


**Figure S15. CD analysis of MEF.** 2 µM **MEF** monomer (thus, 1 µM homodimer) without DNA (blue circles), in the presence of NS DNA (gray circles), and in the presence of E-box containing DNA (orange circles). We see that **MEF** can refold to regain α-helicity after our protein purification scheme. **MEF** undergoes a disorder-to-order transition in the basic region upon the addition of DNA, regardless of specific or nonspecific DNA, which is a common characteristic feature of bHLHZ proteins.

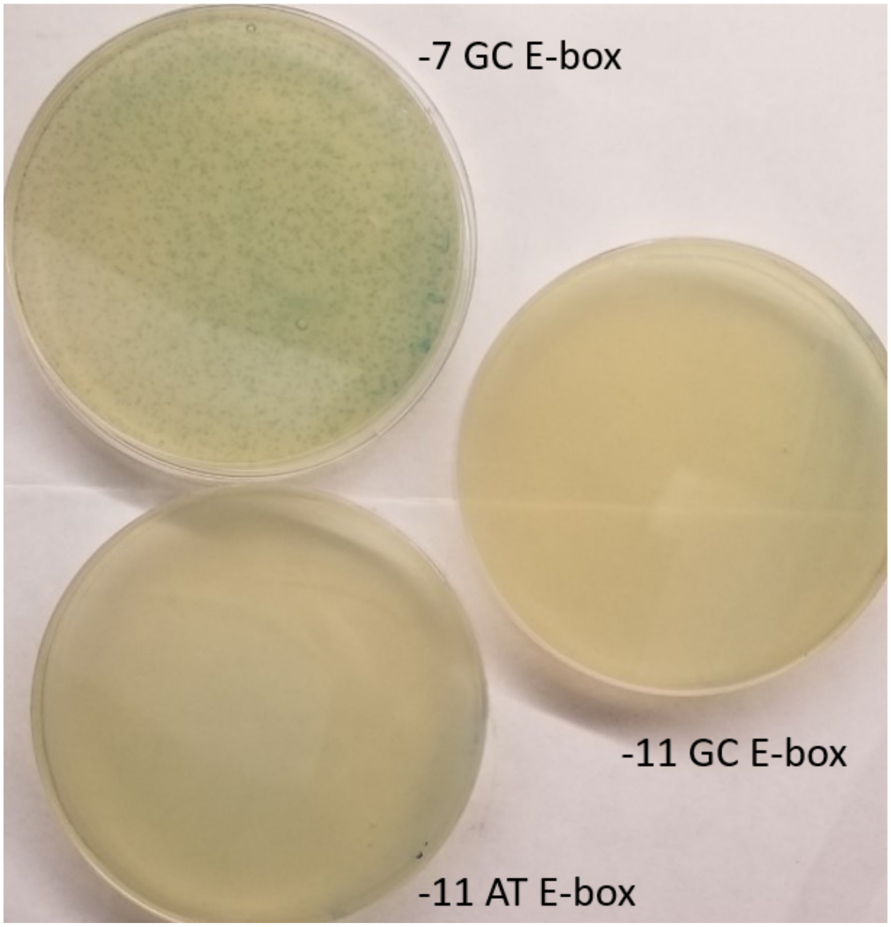


**Figure S16. Plaque assay results for the Accessory Plasmid (AP) library.** 2060 cells harboring the various AP constructs were infected with M13 phage containing SP-ME47 (SP, selection phage; SP-ME47 harbors sequence encoding ME47) to see which AP permits phage propagation. The 2xYT top agar used for the plaque assay was supplemented with Bluo-Gal to allow formation of blue plaques to facilitate visualization of the plaques. Dark blue points are plaques where the 2060 cells with a valid AP construct were successfully infected by the SP-ME47 containing phage, indicating that the AP construct is viable for PACE. Shown here are plaque assays done against SP-ME47 that had been diluted 100-fold to allow for visualization of individual plaques. Only the -7 GC E-box allowed for plaque formation, marking it as the AP to be used in the PACE selection circuit. (Adapted from Popa, Thuronyi, Inamoto, & Shin, manuscript in preparation.)

**Table S1. Protein-DNA complexes. B1H,[a] CD,[b] and $K_d$ values from EMSA.[c]**

| | B1H[a] | | CD, 222 nm[b] | | | EMSA[c] | | |
|---|---|---|---|---|---|---|---|---|
| **Protein** | **E-box** | **AT E-box** | **no DNA** | **E-box** | **NS DNA** | **E-box** | **AT E-box** | **NS DNA** |
| **ME47** | ++++ | +++++ | 23 | 36 | 32 | 15.3 ± 1.6 | 14.8 ± 1.5 | 58 ± 1.0 |
| **ME47/RS** | + | ++ | 7.1 | 13 | 11 | 25.6 ± 9.3 | 29.7 ± 8.7 | — |
| **ME47/RA** | ++++ | +++++ | 29 | 43 | 41 | 20.6 ± 3.7 | 22.3 ± 3.9 | 105 ± 9 |
| **ME47/SC** | +++++ | +++++ | 27 | 40 | 35 | 15.6 ± 1.5 | 16.2 ± 2.1 | 73 ± 18 |
| **ME47/SS** | ++ | ++ | 12 | 21 | 17 | — | — | — |
| **ME47/SA** | ++++ | ++++ | 18 | 30 | 26 | 32.5 ± 4.7 | 28.4 ± 3.6 | 160 ± 7.4 |
| **ME47/CC** | +++++ | +++++ | — | — | — | — | — | — |
| **ME47/AC** | +++++ | +++++ | 27 | 35 | 35 | — | — | — |
| **ME47/AA** | ++++ | ++++ | 26 | 36 | 33 | 27.7 ± 1.7 | 28.5 ± 2.3 | 87.5 ± 5.1 |
| **MEF -1** | ++ | — | — | — | — | 59 ± 5 | — | 111 ± 18 |
| **MEF** | +++++ | — | 17 | 25 | 27 | 7.8 ± 0.6 | — | 96 ± 22 |
| **MEF/LA** | +++ | — | 19 | 33 | 33 | 10.7 ± 1.5 | — | 44.5 ± 6 |
| **MEF +1** | +++ | — | — | — | — | 30 ± 4 | — | 65 ± 7 |

**Additional studies using the AT E-box.** [a] Levels of *HIS3* activity leading to growth relative to growth observed from US0 cells transformed with plasmid encoding specified protein and E-box reporter system. [b] Each percent helicity value is the average of two CD scans of the same sample. Percent helicities determined as described in the experimental section; measurement of protein α-helicity at 222 nm. [c] Each $K_d$ value is the average value from two independent EMSA experiments.
